# The hierarchical structure of canine cognition: two domains and a general cognitive factor

**DOI:** 10.1101/2023.02.07.525704

**Authors:** Zsófia Bognár, Borbála Turcsán, Tamás Faragó, Dóra Szabó, Ivaylo Borislavov Iotchev, Enikő Kubinyi

## Abstract

The complex human environment results in a hard-to-bridge gap between human and animal studies on general cognitive abilities (*g*; colloquially often referred to as “intelligence”). Pet dogs are adapted to our environment, but a convincing demonstration of *g* is missing. Exploratory and confirmatory factor analyses on seven tasks revealed a hierarchical structure with two cognitive domains (problem solving, learning) and an overarching *canine g*. Age did not affect the structure. *Canine g* was negatively linked to age, height, and the owner’s emotional attitude but positively to training level, activity/excitability trait, experienced trauma and normal body condition. The longitudinal analyses over three years differentiated three main trajectory clusters: declining, stable, improving. The diversity of ageing phenotypes supports dogs’ translational value in *g* factor and ageing research.

**One-Sentence Summary:** The *canine g* factor is linked to age, height, training level, personality, body condition, and owner’s attitude.

## Main Text

Intelligence is a topic that has stirred passionate controversies like no other in the social sciences. There is, however, little doubt about the predictive power of IQ (intelligence quotient) scores of humans for important long-term outcomes like career growth, academic success, and social adjustment (*1*). There are still disagreements about what intelligence tests measure (*2*, *3*), but the positive manifold (*4*) is a highly replicable construct (*5*). It emerges independently from the composition of cognitive test batteries (*2*, *6*), and the domain-independent performance is heritable in humans and mice (*6*, *7*). A hypothesized unifying factor behind the positive manifold is the so-called general cognitive factor (or general mental ability, abbreviated as *g*), but to what extent *g* is a single entity is occasionally contested (*2*, *3*). An important and promising intermediate model views *g* as a hierarchical construct (*8*, *9*).

The evolution of general mental ability and how it is affected by the physical versus the social environment are still open questions, requiring comparative efforts (*2*, *10*, *11*). The proof of non-human *g* analogues is currently most solid in rodents (*12*), although among all animals, dogs are the closest to humans in terms of adaptation to the human-made environment and social complexity (*13*). The social domain is the most pertinent of these adaptations for comparative work on *g* factor (*12*, *14*). For instance, a complex social environment is hypothesized to underlie the convergent problem-solving abilities of primates and corvids (*14*). However, very few steps have been taken to realize dogs’ potential for general cognitive ability study (*10*) and no conclusive evidence has yet been provided for the existence of a *canine g* factor. Moreover, previous research on dog cognitive ability had a limited heterogeneity of subjects as well as cognitive tasks, and did not investigate possible affecting factors, including the effect of age (*15*–*17*).

In its alleged role as the underlying factor behind all cognitive faculties (*4*), *g* may help explain how encompassing loss of cognition, as observed in dementia, becomes a risk for some ageing phenotypes. In humans, observations on the relationship between age and intellectual abilities are contradictory, as age-related decline (*18*), stability (*19*), and a mixture of declining and improving capacities have also been reported (*20*). However, for studying individual differences in the ageing curve of cognition, we need longitudinal research (*21*). With such studies, we can identify factors affecting the differences in the onset and the trajectory of age-related decline and disentangle the effects of age and inherent *g* factor differences on the overall performance. Dogs allow us to conduct relevant longitudinal work faster than in humans.

## Results

Here we assessed the cognitive performance of domestic dogs three times, with 1.5 years between measurements and found that a) assessing a diverse sample of dogs with a wide variety of cognitive tasks results in the identification of a single higher-order cognitive factor similar in structure and content to the human *g*; b) as in humans, this *canine g* declines with the animals’ age, and also covaries with the animal’s height, training level, personality, trauma experience, body condition, and the owner’s attitude to keeping dogs; c) the dogs’ health status mediates the age-general cognitive ability relation, and d) the *canine g* enables us to distinguish ageing phenotypes in the dog.

### *Calculating the* canine g

We tested 129 pet dogs of various breeds (age: 3-15 years, weight: 7-45 kg, height: 20-70 cm, except 1 giant dog; Data S1) in a test battery that included seven tasks measuring various cognitive abilities: 1) following human pointing gesture, 2) persistency in object manipulation, 3) one-trial learning in clicker game, 4) success in problem-solving, 5) sustained attention toward moving object/human, 6) associative learning when training for eye contact with a human and 7) short-term spatial memory (see Supplementary Text for details). We used a sub-sample of dogs (N = 32) that participated in the battery twice (average interval: 4.4 months) to assess the re-test reliability of the tasks. We retained six of these tasks for subsequent analyses based on task reliability (ICC > 0.5, Table S2). We subjected these tasks to exploratory factor analysis (EFA) to describe their correlation structure. Unrotated EFA showed that five tasks loaded significantly (> 0.3) on the first factor (Table S4, S5), which we interpreted as the *canine g*. We found that the performances on these five tasks form a positive manifold. A separate rotated EFA on these five tasks identified two domain-specific latent factors (Fig. 1A, Table S6). The first factor was related to individual problem solving and included tasks assessing persistency, speed, and success in finding hidden food. The second factor included tasks that reflect (associative) learning ability. The *canine g* explained 42.8% of the total test variance, which is close to the results obtained with humans and mice (*22*, *23*) and higher than what was found in a previous report on dogs [17% (*15*)].

**Fig. 1.**
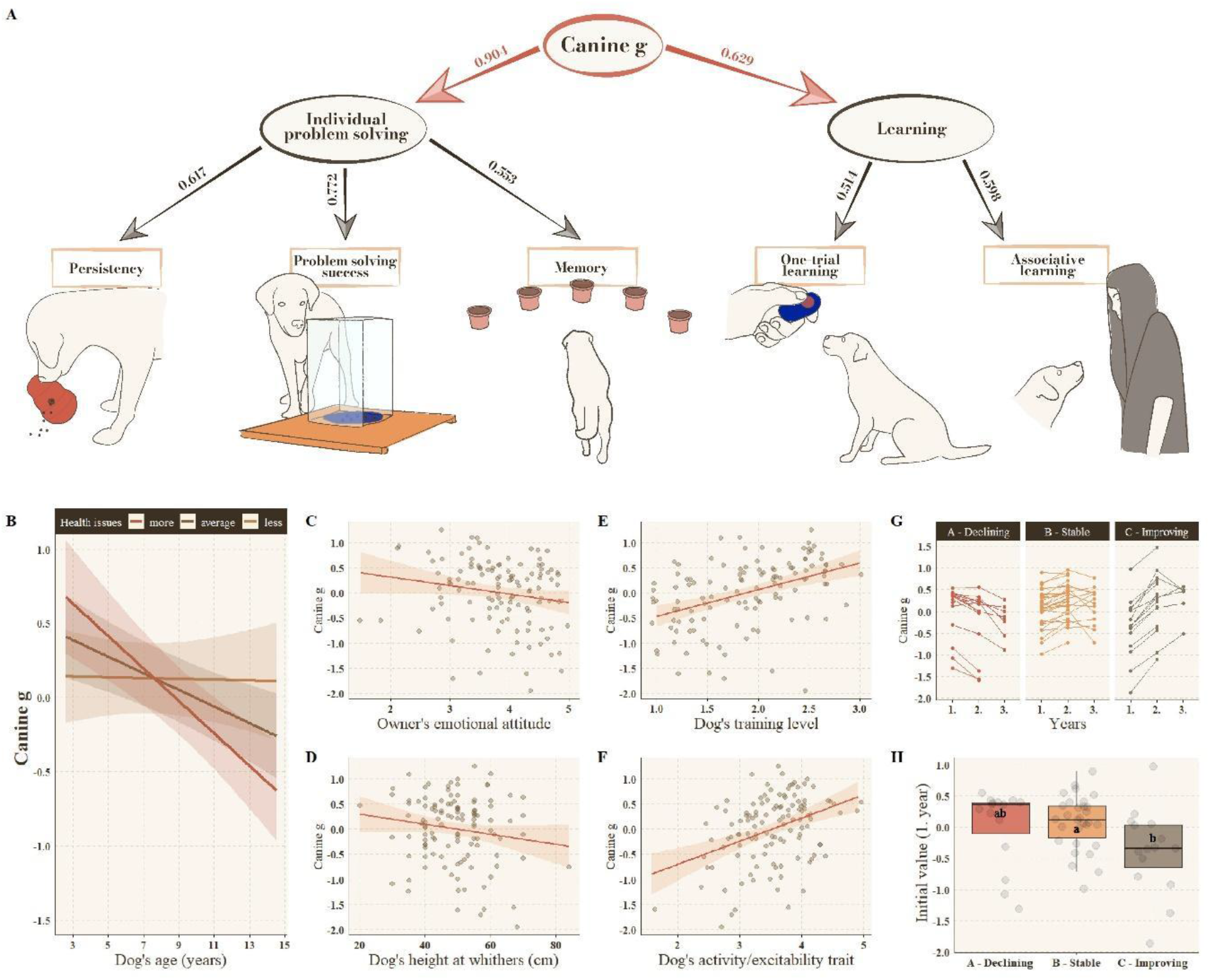
Confirmatory factor analysis best model fit for a *canine g* factor (A), the main effects of different covariates on *canine g* (B-F), and the different trajectories of the changes over time (G-H). (**A**) The performance in the five tasks was entered in the model as observed variables (indicators) and are represented by rectangles (1: Dog manipulates persistently an object to reach the food; 2: Dog quickly solves a problem box puzzle; 3: Dog quickly solves a memory task; 4: Dog shows persistently the previously rewarded behavior 5: Dog quickly learns to form eye contact with a human). Ovals represent latent factors. Arrows from oval to rectangle indicate regression, and values associated with each path are standardized regression coefficient weights. The cross-sectional analyses revealed that (**B**) having health issues strengthened the negative association between the *canine g* and age (orange line: −1 SD, brown line: mean value, red line: +1 SD of health issues). The *canine g* was also negatively associated with (**C**) the owner’s emotional attitude and (**D**) the dog’s height at withers. The *canine g* was positively associated with (**E**) the dog’s training level and (**F**) the dog’s activity/excitability trait. The fitted lines show the estimated main effects. Shaded areas represent 95% confidence intervals around the fits. The dots represent the individual values as partial residuals. The longitudinal analyses differentiated (**G**) three main trajectory clusters: *Declining* (A), *Stable* (B) and *Improving* (C). (**H**) Dogs with lower initial *canine g* factor scores had a higher probability of belonging to the *Improving* than the *Stable* cluster.

Next, we used confirmatory factor analysis (CFA) to explain the origins of the correlations among the observations and characterize the structure of the cognitive abilities. In this analysis, we explored three alternative structures to test the assumptions regarding the existence of a general cognitive factor (*canine g*) and the presence of separate cognitive domains. The structure with the best model fit (Fig. 1A) complied with a hierarchical model (*8*, *9*) that confirmed the presence of both the two cognitive domains (individual problem solving, learning) and an overarching *canine g*.

### *Validation of the* canine g

We next sought to validate the *canine g* with external measurements. The dogs also participated in tests assessing two known covariates of *g*: exploration tendencies and neophilia, which correlated positively with general learning abilities in mice (*24*) and are also conceptually close to the human personality trait Openness to experience, which is a known correlate of IQ in humans (*25*). Moreover, a subset of dogs (N = 60) participated in an independent measure of cognitive performance (discrimination and reversal learning) to which *g* is theoretically related (*26*, *27*), approx. two weeks after they participated in the cognitive battery. Our results complied with both expectations: the *canine g* was found to be positively related to exploration and neophilia (N = 129, r = 0.336, p < 0.001; r = 0.360, p < 0.001, respectively), and negatively to the number of trials required to learn the initial discrimination and the reversal protocol (N = 60, r = −0.418, p = 0.001; r = −0.337, p = 0.009, respectively). These results confirm that the extracted *canine g* represents a domain-general cognitive factor similar both in content and structure to the *g* factor of other species, including humans.

### Controlling for age

In our sample, 41% of the subjects were ≥ 10 years old, which prompted us to control against the possibility that age, specifically cognitive ageing, underlies the emergence of a positive manifold. Cognitive ageing, like *g*, can impact task performances across various domains. Thus, it is essential to exclude that age and *g* are alternative explanations for the observed individual differences in our sample. Replicating all analyses on the age-residuals of the data (see Supplementary Text and Table S4-S6) led to the same results, which supports the independent contributions of *g* and age to individual differences in performance. The *canine g* factor score, like in humans (*18*), correlated negatively with the animals’ age (Winsorized robust correlation: r_w_[CI_95_%] = −0.39[−0.53 - −0.23], t(126) = −4.75, p < 0.001; Fig. S3).

### Exploration of possible modifying effects

In subsequent analyses, both on cross-sectional and longitudinal data, we explored the modifying effects of owner-reported individual features (different demographic and keeping conditions, personality and behavior scales) and owner attitude on the negative association between *g* and age (see the full list of factors in Table S8). Using General Linear Models (Table S9-S15), the cross-sectional analysis showed that overall health moderated the negative relationship between ageing and *canine g:* the association was stronger for dogs with worse health status (ß: −0.341; p = 0.001; Fig. 1B). Additionally, several main effects show direct associations with *canine g* (Fig. S4, see statistical details in Table S15). Independently from age, owners’ general emotional attitude towards dogs was negatively associated with *canine g*: dogs with owners having a more emotional attitude to keeping pets (e.g., anthropomorphizing, see Table S8 for details) had lower *g* scores (ß: −0.197; p = 0.039; Fig. 1C). The dog’s height at withers was also in a negative association with *canine g* (ß: −0.234; p = 0.011; Fig. 1D). Third, the *canine g* was positively associated with the dog’s training level (ß: 0.377; p <0.001; Fig. 1E), and activity/excitability trait (ß: 0.402; p <0.001; Fig. 1F): more trained, more active, and easier to excite dogs had higher *g* scores. Furthermore, the *canine g* factor scores were also generally higher in dogs that had experienced at least one traumatic event according to their owners (ß: 0.566; p = 0.003; Fig. S5). We also found slightly higher *g* scores in dogs with normal body condition than in under- and overweight dogs (ß: 0.493; p = 0.021; ß: 0.637; p = 0.030, respectively; Fig. S6).

In the longitudinal analysis, we examined how the *canine g* changes over three years. Sixty-six dogs attended a second measurement, thirty-one dogs a third, with 1.47 ± 0.39 years between measurements. Using Heterogeneous Linear Mixed Models, we identified different ageing phenotypes, defined by how *canine g* changed relative to the first measurement (Fig. S7; Table S16). We observed three clusters/phenotypes containing 90.9% of dogs who participated on at least two occasions: *Declining* cluster A (22.7%): decline across measurements; *Stable* cluster B (45.5%): relative stability across measurements and *Improving* cluster C (22.7%): improve across measurements (Fig. 1G). The remaining 9.1% of the dogs belonged to three rare clusters and were excluded from further analysis (Table S16). We examined a set of possible predictors (Table S17) for the probability of a dog belonging to one of these three clusters using Fit Multinomial Log-linear Models. Only clusters B and C differed from each other, and only in their initial *g* factor score. Dogs with lower initial *canine g* factor scores had a higher probability of belonging to the *Improving* than the *Stable* cluster. (p < 0.001; A: 0.02 ± 0.61, B: 0.07 ± 0.43, C: −0.39 ± 0.68; Fig. 1H).

## Discussion

Our results consist of findings with both translational and veterinary relevance. The findings support the existence of a *canine g* factor and opened the way for a relatively novel model animal with a high translational value in the study of cognitive ageing. We found that dogs’ general cognitive ability is negatively associated with age and the *canine g*’s malleability across time is linked to health, similar to humans (*18*, *27*). Health problems cannot be completely isolated from lifestyle factors that are known to affect cognitive ageing. Pre-existing or age-related decline in health could lead to lower levels of physical and mental exercise and social engagement, and the attenuation of these preventive factors could be what accelerates the rate of cognitive decline (*28*).

Guidelines for canine welfare can emerge from the observation that inadequate nutrition and emotionally driven keeping practices negatively impact dogs’ overall cognitive capacity. The finding that obese and underweight dogs have lower *canine g* scores than dogs with normal body condition fits well with human results, as both obese and underweight persons perform at a lower level than do people within the normal weight range [see reviewed in (*29*)]. Although the direction of the causality remains to be clarified, these results are most often explained by a common underlying factor being responsible for both weight issues and cognitive function. In the case of dogs, such a common factor can be the owner’s attitude to dog keeping. Indeed, the negative link between anthropomorphizing the animal and its *g* score suggests that this is not a healthy attitude to keeping dogs because it impairs the animals’ cognitive performance.

The finding that higher dogs have lower *g* is the opposite of the human literature. In humans, positive IQ–height correlation was found, which can be due to both genetic and environmental factors (*30*–*33*). Postnatal growth rate was also found to be in positive association with cognitive abilities in humans (*34*). The reasons of the correlation between cognitive ability and height, as well as growth rate are still a matter of debate. As dogs differ in body size substantially, and breed-specific growth velocity also exists [with larger dogs usually having higher growth rates (*35*)], they can be also good subjects of studies revealing the possible connections between height and general cognitive ability.

We found positive associations between the *canine g* and the dogs’ activity, exploration and neophilia both via behavior test and owner reported questionnaire assessment. These behaviors are the closest to the openness to experience personality trait in humans, and openness (sometimes also called intellect) has the strongest link to IQ among the Big Five traits (*25*, *36*). Dogs which are more active and open to new experiences may be easier to train. Dogs’ training level was found to be in a positive correlation with their *g* factor score, which was expected (*37*, *38*) and is in harmony with the human literature on education and IQ (*39*).

The openness to experience trait is also positively associated with successful ageing in humans (*40*, *41*), and there is also some evidence for a three-way, mediation-based relationship between IQ, openness, and the rates of age-related cognitive decline (*41*). However, these interactions are difficult to disentangle in humans, even with complex statistical models, thus, this topic is more amenable to future dog research.

However, the result that trauma (based on owner reports) is associated with higher *g* scores in dogs, contradicts human literature, where trauma is often negatively linked to IQ (*42*–*44*). In humans, secondary trauma, such as fear for loved ones, may contribute to higher IQ (*45*). Thus, one critical question concerns the comparability of what trauma means for dogs and humans. Currently, the findings described here and in the human literature converge on the involvement of traumatic experiences in shaping the IQ/*g* phenotype. However, a better understanding of the proximate mechanisms behind the IQ – trauma association, as well as a deeper understanding of how dogs process such experiences, will be needed to make sense of the counterintuitive direction of the effect we observe in dogs.

In the longitudinal analysis, we aimed to separate our sample into clusters of different ageing trajectories for *canine g*. Contrary to the results of the cross-sectional analysis, we did not find a uniform decline in the *g* score over measurements, in fact, we observed as many dogs to improve their cognitive performance as to decline. Better performance could be due to prior test experience, which we could not separate from a possible systematic cognitive improvement. Importantly, in this study, only animals without major health problems, sensory impairments or conditions negatively affecting their mobility were enrolled. Therefore, we limited our chances to detect cognitive decline to cases where deterioration in cognitive skills was not accompanied by observable health and sensory impairments. Nevertheless, the clusters we described here suggest that cognitive capacity might be crucial for distinguishing ageing phenotypes. Dogs are among the few model species naturally prone to cognitive dysfunction (canine dementia) (*46*), and their risk varies individually, like in humans. Further investigations on larger samples will need to identify factors associated with fast declining *g*. Understanding how these factors act on across the lifespan can enable interventions that allow for sustained cognitive performance in the face of age-related changes for the dog.

## Supporting information

Protocol video

## Acknowledgments

The authors thank the support of the dog owners and Lisa Wallis, Patricia Piotti, Anna Egerer, Alexandra Deés, Bianka Stiegmann, Felícia Erdélyi, Vivien Hemző, Sarolta Marosi and Renáta Böröczki for their help with the data collection; and Andrea Sommese, Soufiane Bel Rhali and Róza Haraszti for their help with inter-observer coding; Péter Ujma, Máté Nagy and Kauê M. Costa for critical feedback and ideas.

## Funding

ÚNKP-20-3, ÚNKP-21-3 and ÚNKP-22-3 New National Excellence Program of the Ministry for Innovation and Technology from the source of the National Research, Development and Innovation Fund ÚNKP-20-3-I-ELTE-950; ÚNKP-21-3-I-ELTE-595 and ÚNKP-22-3-II-ELTE-577 (ZB)

ÚNKP-20-5, ÚNKP-21-5 and ÚNKP-22-5 New National Excellence Program of the Ministry for Innovation and Technology from the source of the National Research, Development and Innovation Fund ÚNKP-20-5-ELTE-337; ÚNKP-21-5-ELTE-1061 and ÚNKP-22-5-ELTE-475 (TF)

MTA Bolyai Research Scholarship BO/751/20 (TF)

European Research Council (ERC) under the European Union’s Horizon 2020 research and innovation programme 680040 (EK)

Hungarian Academy of Sciences via the MTA-ELTE ‘Lendület/Momentum’ Companion Animal Research Group PH1404/21 (EK)

National Brain Programme 3.0 of the Hungarian Academy of Sciences NAP2022-I-3/2022 (EK)

## Author contributions

Conceptualization: BT, DS, IBI, EK

Methodology: ZB, BT, TF, DS, EK

Data collection: ZB, DS

Visualization: ZB, BT, TF

Funding acquisition: ZB, TF, EK

Supervision: EK

Writing – original draft: ZB, BT, TF, IBI, EK

Writing – review & editing: ZB, BT, TF, IBI, EK

## Competing interests

Authors declare that they have no competing interests.

## Data and materials availability

All data are available in the main text or the supplementary materials.

## Supplementary Materials

Materials and Methods

Supplementary Text

Figs. S1 to S7

Tables S1 to S18

References (*47*–*87*)

Movies S1

Data S1

## Notes

### Competing Interest Statement

The authors have declared no competing interest.

