## Supplementary Materials for "The hierarchical structure of canine cognition: two domains and a general cognitive factor"

**The PDF file includes:**

Materials and Methods and Supplementary Text
Figs. S1 to S7
Tables S1 to S18
Caption for Movie S1
Caption for Data S1

**Other Supplementary Materials for this manuscript include the following:**

Movies S1
Data S1

### Materials and Methods and Supplementary Text

*IDENTIFICATION OF A G FACTOR*

Subjects

N = 129 dogs participated in the cognitive test battery. The dogs' age ranged from 2.61 years to 14.54 years (mean age ± SD = 8.38±3.21 years), 48.8% were males, and 77.5% were neutered. The sample consisted of 59 mixed breed dogs and 70 purebred dogs from 33 different breeds. All dogs were middle-sized, their weight ranged from 7 to 45 kg, with the exception of an 80 kg dog (mean weight ± SD = 21.68 ± 7.86 kg, Data S1).

The dogs were required to be free from overt signs of neurological and other physical health problems (as reported by the owner) to participate in the test battery. Additionally, before the test, the dogs' motor skills were assessed by a qualified physiotherapist and a sensory assessment was also performed to exclude subjects with potential visual and/or acoustic impairment [as suggested by (*47*)]. Only animals without major sensory impairments or conditions negatively affecting their mobility (e.g., made it painful for them to walk around) or otherwise prevented them from solving the tasks/detecting the stimuli in the tasks were enrolled in the study.

Re-test data to calculate the task reliability: After the first test (0.23-0.9 years [mean ± SD: 0.37 ± 0.12 years] later), a subset of dogs (N = 32, mean age: 10.92 years, 56.2% male, 78.1% neutered) participated in the test a second time. The procedure and the tasks' rank order were the same in both test sessions, but the objects used in the Exploration and the toys used in the Novel object recognition tasks differed. These dogs did not participate in the longitudinal assessment of their cognitive performance (see later). See Data S1 for more information.

Behavior tests

The cognitive test battery consisted of 10 tasks in total and took place in two experimental rooms: room 1, measuring 5 x 6 m, and room 2, measuring 3 x 5 m. The tasks were provided in a fixed order for all subjects to standardize any carry-over effects between the tasks. The whole battery took approximately 60 minutes to complete, with a short break (Movie S1).
Seven tasks were designed to cover distinct cognitive abilities. These tasks were adapted or modified from published test batteries, required no pre-training, and were aimed to be repeatable with minimal effect of habituation or learning. These seven tasks were used to analyze the correlational structure underlying the individual differences in dog cognition and extract the *canine g*.

Two tasks (the first two of the battery) evaluated the dogs' activity and exploratory tendencies, and one task aimed to assess the dogs' neophilia. These three tasks were used to validate the *canine g* externally. The individuals' propensity to explore a novel environment and their degree of preference for novelty have been consistently found to be positively related to cognitive performance across a wide range of species [see reviewed in (*48*, *49*), including dogs (*50*)]. Furthermore, these traits have also been directly related to *g* in mice (*24, 51*, *52*). Based on these, we expected that higher exploratory tendencies and neophilia would be positively associated with the *canine g*.

Another way to investigate the external validity of the *canine g* was to analyze its correlation with an independent cognitive measure which is a known correlate of *g* in other species. For this aim, a subset of dogs (N = 60) also participated in a spatial reversal-learning task, on average 13 days after participating in the test battery. This task incorporates associative learning, behavioral flexibility, and behavioral inhibition, which are key components of general mental ability in humans and other animals [reviewed in (*12*)]. Based on this, we expected that better performance in this task would be positively associated with the *canine g*.

Cognitive tasks

*1. Pointing*

*Aim*: To measure the dogs' (hereafter D) ability to follow a momentary human pointing gesture when locating a hidden food reward, and also D's ability to shift from a previously rewarded response by pointing to the same side three times in a row, then to the other side during the subsequent three trials.

*Procedure*: The task was conducted in room 2. In the first phase (*Warm-up trial)*, we aimed to ensure that D was comfortable approaching and eating from the containers (pots) used in the test trials. Upon entering the room, the owner (hereafter O) sat down on the chair at the starting position (3 m from the experimenter's (hereafter E) position), took off the leash, and held D by the collar. E called D's attention by saying, "D name + look", showed the treat, then dropped it into the pot and put it in front of her on the ground. Then D was released and allowed to take the food. O and E were allowed to encourage D if necessary.

In the second phase *(Test trials)*, E carried out six test trials. Each started with O sitting in his/her chair, holding D by the collar at the starting position, and E standing 3 m from them, holding two identical pots folded into each other. E called D's attention, and after establishing eye contact, she showed the treat and dropped it into the upper pot. She then shuffled the two pots 2-3 times so that D could not know which one contained the food. Next, she placed them on the marks on the floor to her left and right (the distance between the two pots was 1.5 m). She then called D's attention again ("D name + look") and performed a momentary distal pointing gesture (3 seconds) to the baited pot. After E returned to her starting position (both hands held in front of her chest), O released D. After D made a choice (its nose came within 10 cm of the pot), E removed the other pot before D had the chance to investigate it. If D made a correct choice, it was allowed to eat the food. If D made an incorrect choice, the baited pot was only shown to D. At the end of the trial, O called or led D back to the start position, and the next trial began. There were six test trials, with the first three trials on the same side and the second three on the other side. The location of the first baited pot was counterbalanced among dogs.

*2. Manipulative persistency*

*Aim*: To measure D's willingness and ability to obtain treats from an interactive toy and its persistence in trying to obtain an inaccessible food reward.

*Procedure*: The task was conducted in room 2. In the first phase *(Solvable trial)*, E showed the toy to D (Kong Wobbler™, small or large, depending on D's size), then baited the toy in front of D with 20 pieces of small-sized treats, which could be retrieved by manipulating the toy. The toy was then placed in the middle of the room, and D had 60 seconds to manipulate it while E and O remained in the same positions as in the previous test. If D lost interest, O was allowed to encourage verbally and via pointing at the toy without leaving his/her chair.

In the second phase *(Unsolvable trial)*, the test was repeated with the same procedure, but this time, E baited the toy with a large treat that D could not obtain.

After this test, there was a break (5-10 minutes) during which D remained outside of the test rooms. Then, room 1 was rearranged, removing all the objects.

*3. Clicker game*

*Aim*: To measure D's associative learning ability and behavioral flexibility, that is, its ability and willingness to offer novel behaviors to E in a positive reinforcement setup.

*Procedure*: The task was conducted in room 1. O sat on the chair next to the door and was asked not to communicate or interact with D during the test. E stood in the center of the room with a food pouch filled with sausages on her belt and holding a sound-making device (similar to a clicker but displaying a different sound). She called D to her, then asked it to sit. Once D sat in front of her, she clicked and threw a piece of sausage on the floor. After that, E remained motionless but clicked and rewarded D for presenting any novel behavior, both object- and body-related. If D presented the same behavior repeatedly, E waited until a new behavior was offered while also encouraging D by smiling and nodding but without speaking. The test lasted for two minutes, measured from the first click sound.

*4. Problem-solving*

*Aim*: To assess D's individual problem solving when locating a hidden food reward in a problem box, together with D's behavioral inhibition and flexibility, by systematically shifting the visibility of the food reward and the location of the opening on a problem box.

*Setup*: The task was conducted in room 1. The apparatus had the following dimensions: a 62.5 cm x 53 cm platform with a 22.5 cm x 22.5 cm x 38 cm rectangle box (opaque or transparent) attached to it. The box was closed on the top, bottom, and three sides, with only one side left open.

*Procedure*: In all trials, O sat on a chair 1.5 m from the apparatus, held D by the collar until E baited the apparatus and returned to her starting position next to O. Then O let D free, and D had 45 seconds to obtain the reward. O was allowed to encourage D verbally and via pointing at the apparatus without leaving his/her chair. If D did not succeed in a given trial, E provided the minimal necessary help to D to get the reward to prevent loss of motivation.

In the first phase *(Opaque, trials 1-3),* the apparatus was opaque (wood), D could see the baiting process, and the opening was always in the middle position, facing away from D.

In the second phase *(Transparent, trials 4-10),* the apparatus was transparent (plexiglass) (so D was able to see the food reward inside). In these trials, E prevented D from seeing the baiting process (i.e., the location of the opening on the apparatus) via a visual barrier. The location of the opening was on the same side (left or right) in trials 4 to 6, shifted to the opposite side (right or left) in trials 7 to 9, then shifted to the middle position (facing away from D) in trial 10. The location of the opening in trial 4 was counterbalanced among subjects.

*5. Attention*

*Aim*: To measure D's attentional capture and sustained attention in social and non-social contexts.

*Setup*: The task was conducted in room 1. O sat on a chair approximately 4.5 m from the wall where the stimuli were presented. O was told to ignore both D and the actions of E. D was positioned next to O at the beginning of each context and remained leashed during the entire test.

*Procedure*: The order of the two trials was counterbalanced across subjects. In the *non-social trial (flying object),* E remotely manipulated a yellow plastic frisbee from outside the room by pulling a fishing line through a metal loop in the ceiling in the testing room. The object moved up and down (seemingly on its own) next to the wall facing D for one minute.

In the Social trial *("painting" the wall),* E entered the testing room, silently walked to the wall, and, with her back to D, made up-down movements (as if painting the wall). After one minute, E left the room without looking at D.

*6. Training for eye contact*

*Aim*: To measure D's associative learning ability to sustain social attention, that is, to learn the association between establishing eye contact with E and food reward, and then sustain eye contact with E for increasing durations.

*Procedure*: The task was conducted in room 1. In the first phase *(Training),* D was unleashed, O sat on a chair next to the door and was told to ignore D. E stood in the center of the room holding a clicker-like device in one hand. Both hands were positioned in a relaxed posture by her sides. E had a food pouch on her belt, positioned at her back. First, E called D's attention and threw a piece of food on the floor. Then she remained motionless, and whenever D established eye contact with her, E clicked and tossed a piece of food on the floor. This phase lasted for 20 eye contacts or a maximum of 5 minutes.

There were two conditions in the second phase (*Sustained eye contact*): silent and with distraction, and the order was counterbalanced among dogs. In both conditions, D had to keep eye contact with E for gradually increasing durations to get rewarded. Each condition had five levels: 2 sec, 5 sec, 10 sec, 20 sec, 40 sec. Once D has successfully passed a level, the latency between establishing eye contact with E and the click + reward was increased to the next level. D had three attempts to pass each level. If D failed all three attempts, the test was terminated. In the "with distraction" condition, white noise was played in the background (the mean level of the sound was 49 dB). D received three eye contact retraining trials between the two conditions to maintain motivation.

*7. Memory*

*Aim*: To measure D's visuo-spatial memory.

*Setup*: The task was conducted in room 1. There were five identical pots on the floor, positioned at an equal distance (3 m) from the starting position of D, each pot 1.6 m from the other in a semi-circular arrangement.

*Procedure*: E, O, and D entered the room, O and D walked to the starting position. E called D's attention, showed a piece of food, walked to a pre-selected pot in a straight line from the start, and put the reward in the pot. Then E, O, and D left the room, and outside O distracted D by giving simple commands or petting and talking to D. After 30 seconds, E, O, and D re-entered the room, went to the starting position, and O released D. The trial ended when D found the treat. There were five trials in total, and each container was baited once in a pre-defined order. The order of the baited locations was counterbalanced across subjects.

*Tests used for validating the* canine g

*Exploration and Box rustle*

These two tasks aimed to measure D's activity and exploration in a room containing a wide range of objects. Both were conducted prior to the cognitive tasks, so the individuals were unfamiliar with the test room, and their exploration pattern was not impacted by intervening experience in other tasks of the battery.

*Setup*: Both tasks were conducted in room 1, which included 16 objects to explore. A chair was placed next to the wall, and four larger objects placed in the corners (a large cardboard box with a pink plastic bowl filled with plastic bags placed on top; a small table with a basket filled with plastic bags placed on top; a bedsheet with a small cardboard box on top filled with shredded paper; a waste bin filled with shredded paper) were present in all setups. The other 11 objects were selected from a pool of small-sized everyday objects with different colors and materials. These were placed in a circle around the middle of the room and along the walls.

*Exploration test:* O and D on a leash entered the room together. In the first (*Leashed*) phase of the test, O stood for 20 seconds near the closed door without interacting with the leashed D. In the second (*Unleashed*) phase, O took off the leash on E's signal from outside and released D, ignoring D after that. For 120 seconds, D was free to explore the room while O remained standing next to the closed door. If necessary, O could give a release command to D at the beginning of this test.

*Box rustle test:* O moved slowly around the room, searching for four metal coins, one hidden in each of the four containers placed in the corners of the room. O was instructed to visit the locations in a fixed order (from right to left), spend at least 10 seconds at each location, and ignore D. D was free to move around. The total duration of this task was 60 seconds, and E's signal from outside marked the end.

After this test, E entered the room and greeted the dog-owner pair. She also petted and played with D to ensure that D was not afraid of her and would be familiar with her presence in the subsequent cognitive tasks.

*Novel object recognition*

This task was carried out at the end of the test battery, just before the Memory task. It aimed to assess D's reaction towards and potential preference for novel and familiar toys.

*Setup*: The task was conducted in room 2. The toys used in the test were selected from a pool of six dog toys, and their combination was counterbalanced between dogs.

*Procedure*: In the first phase (*Passive familiarization*), E, O, and D entered the room, O sat down on a chair, let D free, and afterwards ignored D. There were two identical dog toys on the floor, 2 m apart from each other and 1.2 m from D. D was free to interact with them for 30 seconds.

In the second phase (*Active familiarization*), E approached and interacted with both toys (picked them up individually and engaged D's attention by saying, "what do I have" and "look at this" in a happy voice). This phase also lasted for 30 seconds. After phase 2, O and D left the room for 5 minutes while E switched one of the toys to a new one.

In the third phase (*Test phase*), after re-entering the room, O sat down, released D, and D was free to interact with the toys for 60 seconds.

*Reversal learning task*

This task was carried out 1 to 94 days after the cognitive battery (mean: 13 days). The task was based on the cognitive bias paradigm (*53*). It contained a discrimination phase aiming to assess D's ability to learn the association between the location of a stimulus and the reward. A reversal phase measured D's ability to re-learn the association when the positive and negative stimuli were reversed.

The task took place in room 2. The stimulus was a blue plastic plate (20 cm in diameter), and the discrimination was based on the location of this plate (left or the right-hand side of E). The side used as the positive stimulus (S+) was counterbalanced among the dogs, and when the bowl was placed on this side, it always contained a small piece of food, while on the negative side (S-), the bowl was always empty. D received the positive (S+) and the negative (S-) stimuli in consecutive trials presented in a fixed semi-random order (S+S+S-S+S-S-), which was repeated until the criteria (see below) was reached or for a maximum of 50 trials.

*Procedure:* At the beginning of each trial, O was sitting on a chair approximately 3 m from E and held D by the collar or leash. E turned its back to O and D and baited (or pretended to bait) the plate. Then she turned back and called D's attention ("name + look"). Once she established eye contact with D, she put the plate on its pre-determined location (left or right side, ~1m from E). O was instructed to let D go immediately as the plate touched the floor. If D did not start moving when released, O was allowed to encourage it verbally (e.g., "Go!", "It's yours") or by gently touching it. Apart from this, no other forms of communication were allowed. D had 15 seconds to reach the plate (and, in the case of S+, eat the food), then E picked up the plate, and O called D back for the subsequent trial.

In each trial, E recorded the latency to reach the plate, measured from the moment the plate touched the floor until D was < 15 cm of the plate (defined by markings on the floor). If D did not approach the plate, E gave the maximum latency (15 sec). D was deemed to have learned the association between the stimulus and the food when the longest latency in the last five positive (S+) trials was shorter than any latency in the last five negative (S-) trials. The testing was terminated if D did not reach this criterion within 50 trials or refused to leave the chair's proximity for three consecutive trials. If D passed the criteria of the discrimination phase, the test continued with the reversal phase after a short break. In the reversal phase, the procedure was the same, but the S+ and S- locations were switched, i.e., if the S+ was on the left side, it became the right side. Again, D had a maximum of 50 trials to learn the reversed association, the criteria of learning was the same as in the discrimination phase.

*Variables*

The behavior tests were video-recorded, and the videos were later coded using the Solomon Coder program (beta 19.08.02) and a scoring sheet. In contrast to most human and animal general cognitive ability tests, we did not select a single reference variable *a priori* for each task. Instead, we collected multiple variables for each task (Table S1) and used principal component analysis (PCA) to extract a component representing the dogs' aggregate performance. To assess the inter-observer reliability, we randomly selected a sample of 20 dogs in each task to be coded by a second observer. The inter-observer reliability was calculated using Intraclass correlation coefficient (ICC) for all variables. We found all observations to be reliable (Table S1).

Performance in the cognitive tasks and task reliability

To obtain composite scores reflecting the animal's test aggregate task performance in the cognitive tasks, we ran a PCA with Oblimin rotation for each task separately, except for the Pointing task where only a single variable was coded. We decided upon this method because, in contrast to humans and mice, there are no standardized protocols established to assess particular cognitive abilities yet. Thus, our battery may include tasks where the performance is strongly influenced by non-cognitive factors (i.e., motivation or training experience), tasks where the performance is related to multiple cognitive abilities, or where different abilities are engaged across different subjects. These task impurities can obscure the underlying structure in the individual variance (*2*). Running a PCA for each task was a suitable way to filter out variables whose individual variance had a strong non-cognitive source. Moreover, extracting components that represent the common variance shared across multiple variables could also help reduce measurement error and other variable-specific effects (*54*).

In cases where the variables were categorical (Attention task), the PCA was run based on a polychoric correlation matrix; otherwise, normal correlation matrixes were used. The number of components extracted was decided by parallel analysis in all analyses. Variables that failed to load > 0.5 on any components were removed. We used Kaiser–Meyer–Olkin (KMO) measure and Bartlett's sphericity test to determine the sampling adequacy and Cronbach's alpha coefficient to assess the internal consistency of the items.

We extracted seven components in total (Table S2). The KMO value was > 0.5 for all analyses. In 5 out of 6 task-level analyses, all variables loaded on a single component, suggesting that they all share a common source of variance. In one task (Clicker game), we found two components.

The task reliability was investigated using Intraclass correlation (ICC, two-way mixed model) on the subsample of dogs (N=32) that participated in the battery two times.

The Pointing task was found to have low repeatability (Table S2), which could mean that a higher portion of the individual variance in this task is due to random or measurement error or factors unrelated to cognition (*55*). Thus, this task was excluded from further analyses. The reliability of the other six tasks (seven components) was significant and at least moderate (ICC > 0.5), suggesting that the intra-individual variability was not too high for the majority of the tasks.

Correlation structure among the tasks

To explore and describe the correlation structure among the different tasks, we subjected the seven components to exploratory factor analysis (EFA) using the principal axis factoring method of SPSS (version 28). We conducted the EFA both without rotation and with Oblimin rotation. The former analysis was aimed to identify the psychometric *g* based on the first unrotated factor [see (*56*, *57*)]. The rotated analysis aimed to assess if there are distinct cognitive domains, that is, specific groups of cognitive tasks which share a part of their variance with each other but not with other tasks. In this latter analysis, we removed the cognitive components that failed to load > 0.3 on any factors (*58*).

We also assessed if/how much age affects cognitive structure. Many cognitive functions have been shown to decline with age in dogs (*59*, *60*). Therefore, positive correlations among the performance of various cognitive tasks may be partly attributable to the combined effect of age (*54*). In order to confirm that the extracted factors are not a statistical artifact of this confounding factor, we replicated the EFAs on the age-residuals of the data. The age association of all seven components was investigated using linear and quadratic regression models to obtain the age-residuals.

For all seven components, the age relations were primarily linear (Table S3). Therefore, the residuals were extracted from the linear regression models, and we replicated all analyses on the age-residuals of the data.

The parallel analysis suggested three factors to be retained on both the raw data and age-residuals of the test data. The KMO measure of sampling adequacy was 0.636 and 0.609 on the two datasets, respectively, and Bartlett's test of sphericity was highly significant ($p < 0.001$), confirming that the sample size was adequate for the EFA.

The unrotated EFA were in accordance with the correlation analysis and showed that five components loaded on the first factor, which we termed the *canine g* (Table S4). The results were the same when the analysis was run on the age-residuals of the data. The two components that did not load on this factor significantly (> 0.3) were Flexibility and Attention to object. If these two were removed from the analysis, the first factor (*canine g*) explained 42.8% of the total variance on the raw data and 40.8% on the age-residuals (Table S5).

In the rotated analysis (Table S6), the KMO measure of sampling adequacy was adequate for both datasets (0.695; 0.677, respectively), and Bartlett's test of sphericity was highly significant ($p < 0.001$). The parallel analysis suggested two factors to be retained, and they together explained 63.4% of the total variance on the raw data and 62.3% on the age-residuals.

The five components that compose the *canine g* formed two factors. Performance on three tasks (Manipulative persistency, Problem solving, and Memory) loaded on Factor 1, which we termed as *Individual problem solving*. Higher scores mean higher persistence in searching for hidden food, faster success in obtaining it, and better memory in recalling where it was hidden. The two tasks that split off to form the second factor (Clicker game and Training for eye contact) are related to associative learning; thus, we termed this factor as *Learning*. Higher scores mean faster learning of the association between establishing eye contact with the experimenter or any performed behaviors and a reward. However, both tasks were performed using a clicker-like device; thus, their shared variance might also be previous experience with using a clicker. The two factors correlated positively with each other (raw data, r = 0.466, age-residuals, r = 0.434), suggesting that they share a common source of variance.

Confirmatory factor analysis

While EFA is well-suited to describe and summarize the correlation patterns among the subjected variables, it is not well suited to explain where the correlations among the observations originate from. The variance in all cognitive measures has three separate components: (1) a *g*-specific part common in all measures, (2) a domain-specific part common in all tasks assessing a given ability, and (3) a task-specific (or error) variance unique to that given task (*8*, *61*). EFA uses the total variance for extracting the factors, including domain-specific and unique variance, preventing us from assessing where the correlations originate. EFA will extract a general factor even if there are multiple independent and equally strong causes at work (*62*). Moreover, EFA is a descriptive, formative model, so it is not a reliable instrument for causal inference. The extracted factors are linear combinations of the observed measurements, and as such, the factors are "causally" determined by the observations, not the other way around. Confirmatory Factor Analysis (CFA) is better suited to test that the cognitive factors extracted are causal determinants of the test performance (*63*). CFA uses an inferential model, which can partition the variance of the observations and test whether the hypothesized latent factors indeed account for all the correlations among the observations. CFA tests only correlations indicated in the model and assumes all other relationships to be zero. Thus, it is suitable to test one of Spearman's original assumptions in the *g* theory, that the residual correlations between the observations left after controlling for the common factor should be zero (*4*). The analyses were carried out using the AMOS program (version 27). Posterior predictive p-value (PPP) was used to reject models, where PPP < 0.05 indicates poor model fit. The alternative models were compared using AIC and BIC values and maximum likelihood criteria: $Chi^2$ and RMSEA values, where RMSEA < 0.1 was considered acceptable, and < 0.05 a good model fit (*64*). The CFA analyses were run only on the subjects with no missing data (N = 124 dogs) to ensure that the sample was the same as in the EFA analysis. Similar to the EFA, the CFAs were also conducted both on the raw data and the age-residuals of the data, to assess the effect of age on the cognitive structure.

We fitted three alternative models on the data. The cognitive components were entered in all models as the observed variables (indicators), each with its unique error variance. We posited a various number of higher-order latent factors influencing the components according to the structure of the actual model. The residual covariances (i.e., covariances of the error variances) among the cognitive components, even those between components of the same latent factor, were fixed to zero, which posited that all correlations among the different tasks would be accounted for by their common latent factors.

1) The *Hierarchical g* model (Fig. S1A) assumed the existence of both the separate cognitive domains and also the general cognitive factor (*canine g*). This model included Individual problem solving and Learning as first-order latent factors (cognitive domains), which directly affected the five cognitive components according to the pattern found in the EFA. These factors also had their own error variance representing domain-specific variance. A single higher-order latent factor (*canine g*) was set at the apex, influencing the first-order factors.

2) The *No g* model (Fig. S1B) assumed the existence of separate cognitive domains but not the *canine g*. It included the two first-order latent factors (Individual problem solving and Learning) directly affecting the five cognitive components according to the pattern found in the EFA, but this model assumed that the two first-order latent factors are independent of each other.

3) The *g-only* model (Fig. S1C) assumed the existence of the *canine g*, but not the separate cognitive domains. This model included the five cognitive components and posited a single higher-order factor (*canine g*) directly influencing all of them.

The results showed that in both the raw data and the age-residuals of the test data, the best model fit belonged to the Hierarchical *g* model (Fig. S2). This model version also fitted the data significantly better than any of the other models (smallest $Chi^2$ difference=6.18 with df difference of 1), which confirmed the existence of both the two cognitive domains and the *canine g* itself. For further analyses, we calculated the factor scores of the two cognitive domains as the average of the related task performances and the score for *canine g* as the average of the two cognitive domain scores.

<u>External validation of the extracted factors</u>

*Exploration and novelty seeking*

Next, we aimed to determine if the individual's activity, exploratory tendencies, and preference for novelty would correlate with the dogs' general cognitive performance (*canine g*). The behaviors coded during the Exploration, Box rustle, and Novel object recognition tests were subjected to principal component analyses – similar to the cognitive tasks – to extract an aggregate measure of test performance (Table S7). In the case of the Novel object recognition test, two components emerged, and only the first indicated the dogs' preference for the novel object; we expected a positive association with the *canine g* in the case of this component.

Activity during the Exploration test, as well as the Preference for novelty component correlated significantly and positively with the *canine g* ($r = 0.336$, $p < 0.001$; $r = 0.360$, $p < 0.001$, respectively), and with the Individual problem solving cognitive domain ($r = 0.306$, $p < 0.001$; $r = 0.325$, $p < 0.001$). The Learning domain had only a weak association with these two traits ($r = 0.246$, $p = 0.006$; $r = 0.256$, $p = 0.004$). The performance in the Box rustle test had no significant association with any cognitive factors. Nevertheless, these results suggest that the dogs' general cognitive abilities are positively related to the individuals' propensity for exploration and their preference for novelty, similar to what was found in other species, which provides evidence for the external validity of the *canine g*.

*Reversal learning*

Finally, we also investigated the external validity of the *canine g* extracted from the cognitive battery on a sub-sample of N = 60 subjects by analyzing their correlation with an independent measure of the associative and reversal learning ability of the dogs.

The number of trials required to learn the initial association between the stimuli and reward correlated significantly and moderately with the *canine g* ($r = -0.418$, $p = 0.001$), and weaker but still significant with both cognitive domains (Individual problem solving: r = -0.325, p = 0.011; Learning. r = -0.353, p = 0.006). The number of trials required to learn the reversed association correlated significantly with *canine g* (r = -0.337, p = 0.009) and Individual problem solving (r = -0.334, p = 0.009). For both variables, dogs with higher cognitive performance scores required fewer trials to learn the initial or reversal association, which fits the literature, i.e., learning performance is a known correlate of *g* (*12*). This association provides further evidence that the *canine g* represents a domain-general cognitive factor and shows some similarity in the content to the general cognitive ability factor found in other species, including humans.

*CROSS-SECTIONAL ANALYSIS*

<u>Subjects</u>

The same 129 dogs were included in the cross-sectional analysis as in the identification of the *g* factor.

<u>Questionnaire</u>

To collect basic information regarding the demographic attributes of the dog and the owner as well as the social attributes of their interactions, we used a demographic questionnaire described in (*65*, *66*).

To measure dog personality traits, we used the five factors of the Dog Personality Questionnaire [DPQ (*67*)]. We used 5-point Likert scales instead of 7-point in the original 45 items, but this simplification did not lower the reliability, based on Wallis et al. (2020) report (*66*). The factors were calculated by averaging the items (reversed if necessary) following the original structure published in (*67*).

Finally, the questionnaire included three additional queries, 10 questions about the owners' attitude towards dogs, 30 items about the cognition and communication of the dog, and 11 items asking about the dogs' character. The owners indicated their agreement/disagreement with each statement using a 5-point Likert scale in all these questions. The questionnaires were filled out around the date of the first test occasion (mean ± SD = 0.998 ± 3.61 months; range: -14.95 – 15.15 months).

<u>Data reduction in the questionnaire</u>

As several variables co-varied and thus cannot be considered independent measures, we applied PCA on an extended sample of questionnaire answers (N=1532) to reduce the number of variables. In this PCA analysis, we included 19 various demographic variables and, as they were either nominal or ordinal scales, we used a heterogeneous correlation matrix [“hector” function of “polycor” package (*68*)] as the basis of the PCA. As earlier, the number of potential components was determined by parallel analysis [“fa.parallel” function of “psych” package (*69*)]. Then we cleaned the PC model by step-by-step elimination of single component items and items with low loading (<0.4). From the resulting structure, the first three components [labeled as Training level (standardized Cronbach's alpha: 0.723), Family (standardized Cronbach's alpha: 0.533), and Health issues (standardized Cronbach's alpha: 0.620); see Table S8] had acceptable internal consistency, thus were kept for later analyses. The component scores were calculated as item averages. Variables that have fallen out of the PCA were entered in later models as individual items. Aside from the demographic and personality questionnaire parts, we had three more sub-questionnaires analyzed with separate PCAs.

1) A set of 10 questions about the owners' attitude towards dogs formed two scales based on PCA: emotional attitude (standardized Cronbach's alpha: 0.710) and doggy lifestyle (standardized Cronbach's alpha: 0.660).

2) Cognition and communication-related items (N = 30), originating mainly from (*38*, *70*), formed a three components structure: *Communication* (standardized Cronbach's alpha 0.858), *Signs of decline* (standardized Cronbach's alpha: 0.769), and *Uncontrollability* (standardized Cronbach's alpha: 0.620).

3) Finally, 11 items asked about the dogs' character. The PCA revealed a three-component structure; however, only the first had acceptable internal consistency; thus, we kept only this scale, which we labeled as Sociability/trainability (standardized Cronbach's alpha: 0.720). The setup of these analyses was the same as described above.

<u>Age association of the *canine g*</u>

To check the overall relationship between the dogs' age at the time of the first behavior test and the resulting *g* score we used robust correlation analysis ["ggscatterplot" function of "ggstatsplot" package (*71*)]. We decided to use this approach as it is not sensitive to skewness and kurtosis deviations of the variable, and in both cases, we have significant deviations from normal distribution [*canine g* score - D'Agostino test: skew = -0.701, z = -3.126, p-value = 0.002; Anscombe-Glynn test: kurt = 3.099, z = 0.524, p-value = 0.600; age - D'Agostino test: skew = -0.307, z = -1.473, p-value = 0.141; Anscombe-Glynn test: kurt = 1.907, z = -5.462, p-value < 0.001; "moments" package (*72*)].

The results showed a negative relationship with age (Fig. S3).

In further analyses, we applied power transformation of the *g* score to normalize it. The exponent ($\lambda$=1.5) was identified with Box-Cox analysis ["boxcox" function of "MASS" package (*73*)] and normality was checked (D'Agostino test: skew = -0.245, z = -1.175, p-value = 0.240; Anscombe-Glynn test: kurt = 2.44, z = -1.514, p-value = 0.130).

<u>Stepwise model selection</u>

Next, we explored the modifying effects of owner-reported individual features (demographic and keeping condition, personality, and behavior scales) and owner attitude on the negative association between *canine g* and age.

*Step 1. Separate models*

To avoid overparameterization, first, we used four separate General Linear Models (lm) with backward elimination-based model selection (step function) to find the AIC-based parsimonious models. Each initial model contained a specific set of variables (numbers refer to Table S8) and their interaction with age. All initial models only including main effects were checked for multicollinearity ["check_colinearity" function of "performance" package (*74*)]. All VIFs were under 2.

Model 1 contained individual features somewhat independent from the owner and the keeping conditions: 2) sex; 3) reproductive status; 4) breed (grouped later into purebreds and mixed breeds); 5) weight; 6) height at withers; 7) body condition (underweight, normal, overweight); 16) Health issues; 14) Training level; 9) Previous experienced trauma. For the final model, see Table S9.

Model 2 contained keeping conditions: 10) Time spent playing with the owner per day; 11) Off-leash activity per day; 12) Time spent alone per day; 13) Other dogs in the household; 8) owner's age; 15) Family level. For the final model, see Table S10.

Model 3 contained the DPQ scores: 17) Fearfulness; 18) Aggression towards People; 19) Activity/Excitability; 20) Responsiveness to Training; 21) Aggression towards Animals. For the final model, see Table S11.

Model 4 contained the other questionnaire scales: 22) Owner's emotional attitude; 23) Owner's doggy lifestyle; 24) Communication; 25) Signs of decline; 26) Uncontrollability; 27) Sociability/trainability. For the final model, see Table S12.

*Step 2. Combined model results*

Next, to explore which variables affect the *canine g* and how these modify the age effects, we defined a *Combined model* which included all the significant and trend-level effects and interactions found in the former models, then again ran the AIC-based model selection process to reach the most parsimonious model. This *initial combined model* included*:* dog age, reproductive status, height at withers, body condition, Health issues, Training level, Previous experienced trauma, dog age * reproductive status, dog age * Health issues, dog age * Training level, dog age * Previous experienced trauma, Aggression toward People, Activity/Excitability, Responsiveness to Training, dog age * Aggression toward People, dog age * Responsiveness to Training, Owner's emotional attitude, Owner's doggy lifestyle, Signs of decline, Sociability/trainability, dog age * Owner's doggy lifestyle. Following this automatic model selection, applying a conservative approach, we further eliminated marginally significant and trend effects based on F tests (drop1 function) (Table S13), and applied the Benjamini-Hochberg correction to control for multiple comparisons. The *strict, final model* included: dog age, height at withers, body condition, Health issues, Training level, Previous experienced trauma, Activity/Excitability, Owner's emotional attitude, dog age * Health issues (Fig. S4, S5, Table S14, S15).

A Tukey post-hoc test was used for comparisons between the body condition groups [“emmeans” function of “emmeans” package (*75*)] and Simple slopes analysis [“interactions” package (*76*)] for the dogs’ age and health issues interaction.

These post-hoc tests showed that dogs with normal body condition had slightly higher *canine g* factor scores than under- (ß ± SE: 0.493 ± 0.210; t = 2.344; p = 0.054) and overweight dogs (ß ± SE: 0.637 ± 0.289; t = 2.204; p = 0.075). Under- and overweight dogs did not differ from each other (ß ± SE: 0.144 ± 0.324; t = 0.445; p = 0.897; Fig. S6).

The Simple slopes analysis revealed that the negative association between dogs’ age and their *canine g* factor score was significant only if the dogs’ health issue score was above 0.967. This age association was more negative if the health issue score was higher (at 0.62 (-1 SD) health issue score: b[CI95%] = -0.01[-0.09, 0.08]; Z = -0.14; p = 0.89; at 1.24 (mean) health issue score: b[CI95%] = -0.11[-0.17, -0.05]; Z = -3.69; p < 0.001; at 1.87 (+1 SD) health issue score: b[CI95%] = -0.22[-0.31, -0.13]; Z = -4.75; p < 0.001).

*LONGITUDINAL ANALYSIS*

<u>Subjects</u>

Sixty-six dogs participated in the longitudinal assessments [28 males (7 intact), 38 females (3 intact)]. Thirty-five dogs two times, 31 dogs three times. Between the three occasions, on average there were 1.47 ± 0.39 years. The study covered on average 2.17 ± 0.50 years per dog (1.31 – 2.96 years). Dogs' age at first participation ranged from 2.61 years to 12.3 years (mean age ± SD = 7.98 ± 3.04 years). The sample consisted of 36 mixed breed dogs and 30 purebreds, and their weight ranged from 8 kg to 44 kg (mean weight ± SD = 19.9 ± 8.15 kg). See Data S1 for more information.

Statistical analysis

To analyze individual trajectories of *canine g*'s changes, we used R statistical software [version 3.6.3; (*77*)] in Rstudio (*78*). For making distinct clusters based on dogs' individual trajectories, we used Heterogeneous Linear Mixed Models ["hlme" function of "lcmm" package (*79*)]. Then we analyzed which factors and covariates affect the probability to belong to different clusters using Fit Multinomial Log-linear Models ["multinom" function of "nnet" package (*73*)]. First, we made basic models, which included only one explanatory factor. Then, we used bottom-up model selection for finding the best complex model ("anova" function of "stats" package), where the inclusion criteria were a significant likelihood ratio test for each tested variable.

Results

*Clusters based on individual trajectories*

We could distinguish six clusters based on how *canine g* changes from the first measurement (Fig. S7). The clusters' mean *g* score changes are presented in Table S16. The *g* score of 25.7% of dogs declined, but they belonged to two separate clusters, *cluster A* and *cluster D*. We had to exclude *cluster D* from further analysis due to the low sample size (N = 2). At the first measurement, both dogs in cluster D were 12 years old, and we were not aware of lifestyle changes. The majority of dogs (45.5%) belonged to the relatively stable *cluster B*, where the *g* factor score increased slightly at the second measurement and decreased slightly at the third measurement, thus was equal to the initial *g* score. 28.7% of dogs could improve their performance from the first measurement, but they belonged to three separate clusters, *C*, *E* and *F*. Again, we had to exclude *clusters E* and *F* from further analysis due to the low sample size (N=2, both). The dogs in *cluster E* were 6.3 and 10.7 years old at their first tests. The older dog had a scheduled, routine ultrasonic tartar removal under anesthesia two months before the third measurement (when he was 13 years old). The owner reported that the dog showed mental deterioration since the surgery, and two months after the measurement, the owner opted for euthanasia due to it. In the case of the six-year-old dog, lifestyle change is unknown. The improvement in *cluster F* was twice as much as in *cluster C*. The dogs were both eight years old at the first measurement. Their cognitive performance was low at that time, but they could significantly improve their performance already at the second measurement, one and a half years later. This improvement was probably because their owners did not train their dogs before participation but started to train them and take part in intense activities after the first measurement (personal communication with the owners), achieving a significant improvement.

*Differences between the clusters*

The result of the AIC-based bottom-up model selection (see Table S17) is that only dogs' initial *g* factor score (p = 0.035) affected the probability of belonging to different clusters. The model selection criterium was AIC decreasing by at least two values. According to post-hoc tests, only cluster B differed from cluster C (Table S18).

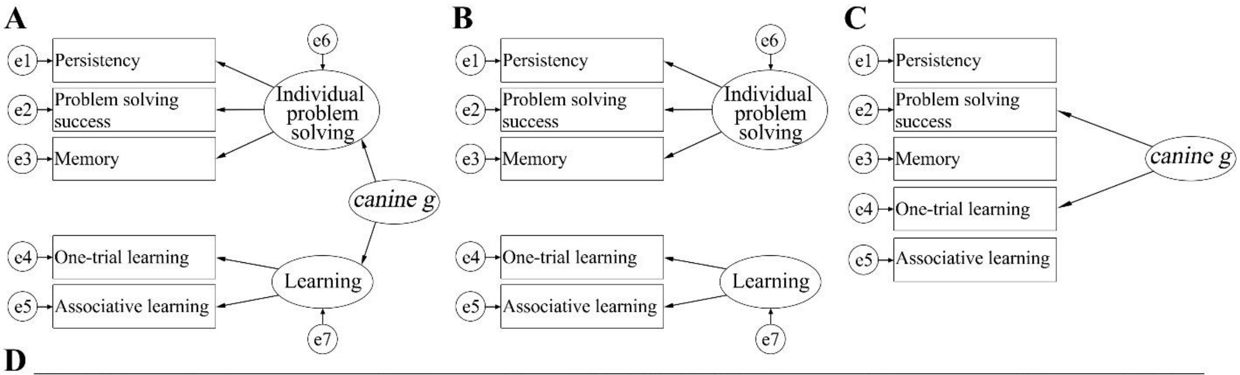


**D**

| Model (raw data) | Chi$^2$ | df | RMSEA (90 % CI) | PPP | AIC | BIC |
|---|---|---|---|---|---|---|
| Hierarchical g model | 4.365 | 4 | 0.027 (0.000-0.141) | 0.510 | 26.365 | 57.388 |
| No g model | 18.533 | 6 | 0.130 (0.066-0.200) | 0.025 | 36.533 | 61.916 |
| g only model | 10.545 | 5 | 0.095 (0.000-0.176) | 0.150 | 30.545 | 58.748 |
| **Model (age-residuals)** | | | | | | |
| Hierarchical g model | 2.812 | 4 | 0.000 (0.000-0.116) | 0.716 | 24.812 | 55.835 |
| No g model | 13.286 | 6 | 0.099 (0.021-0.172) | 0.114 | 31.286 | 56.668 |
| g only model | 8.655 | 5 | 0.077 (0.000-0.161) | 0.250 | 28.655 | 56.858 |

**Fig. S1. Alternative model structures compared in CFA.** (A) Hierarchical *g* model; (B) No *g* model; (C) *g*-only model. (D) Model fit indexes of the three models on both the raw test data and the age-residuals. In all models, the components were entered as observed variables (indicators) and are represented by rectangles. Latent factors are represented by ovals. Arrows from oval to rectangle indicate regression, circles e1 to e7 represent error variance.

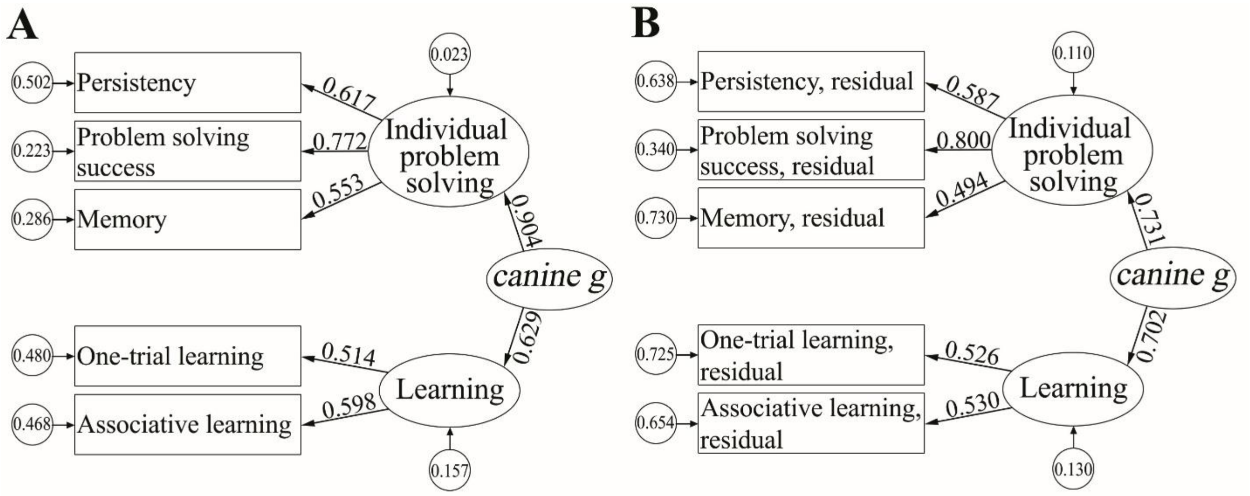


**Fig. S2. The Hierarchical *g* factor model results on the raw test data (A); and the age-residuals of test data (B).** The five cognitive components were entered as observed variables (indicators) and are represented by rectangles. Individual problem solving, Learning, and *canine g* were entered as latent factors and are represented by ovals. Arrows from oval to rectangle indicate regression, and values associated with each path are standardized regression coefficient weights. Circles represent error variance, i.e., the part of the variance unique to that variable/factor.

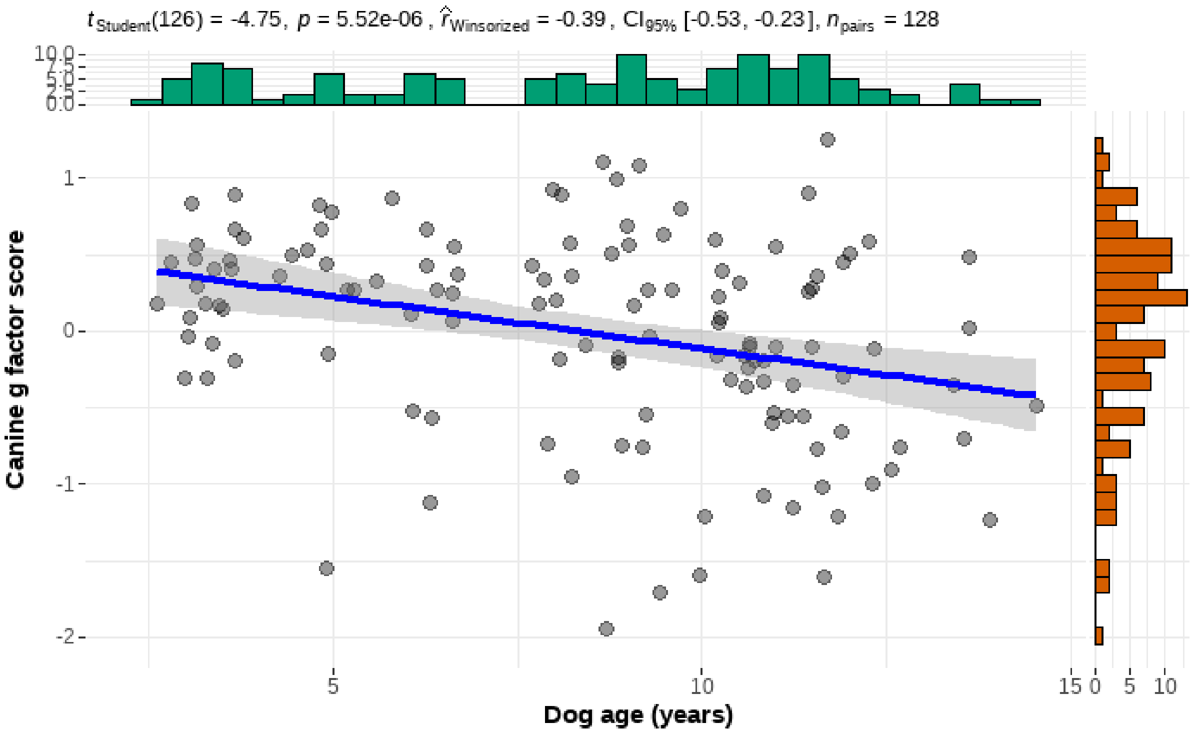


**Fig. S3. The negative correlation between age and *g* score**

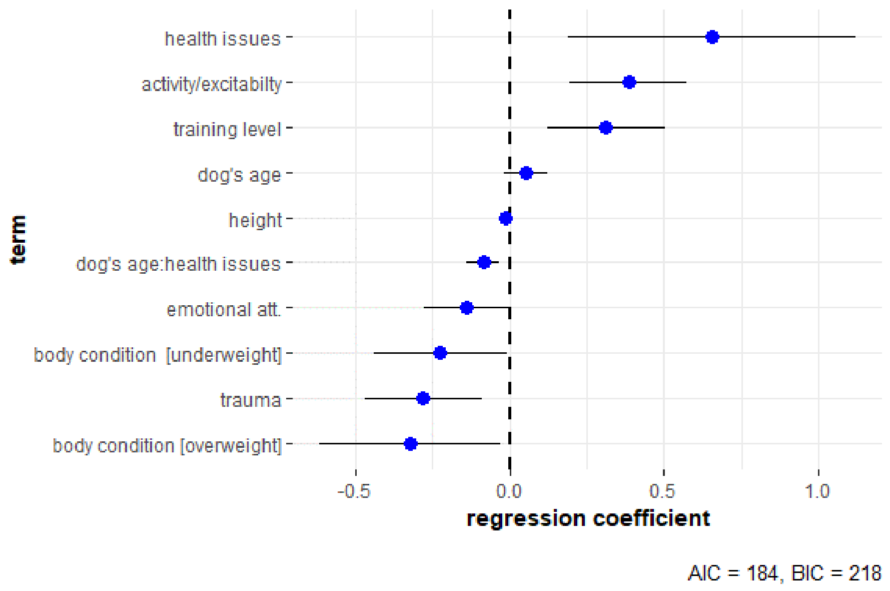


**Fig. S4. Effect plot from the combined model.** The blue dots show the model estimates of the different effects (excluding the intercept), while the whiskers show the 95% confidence intervals

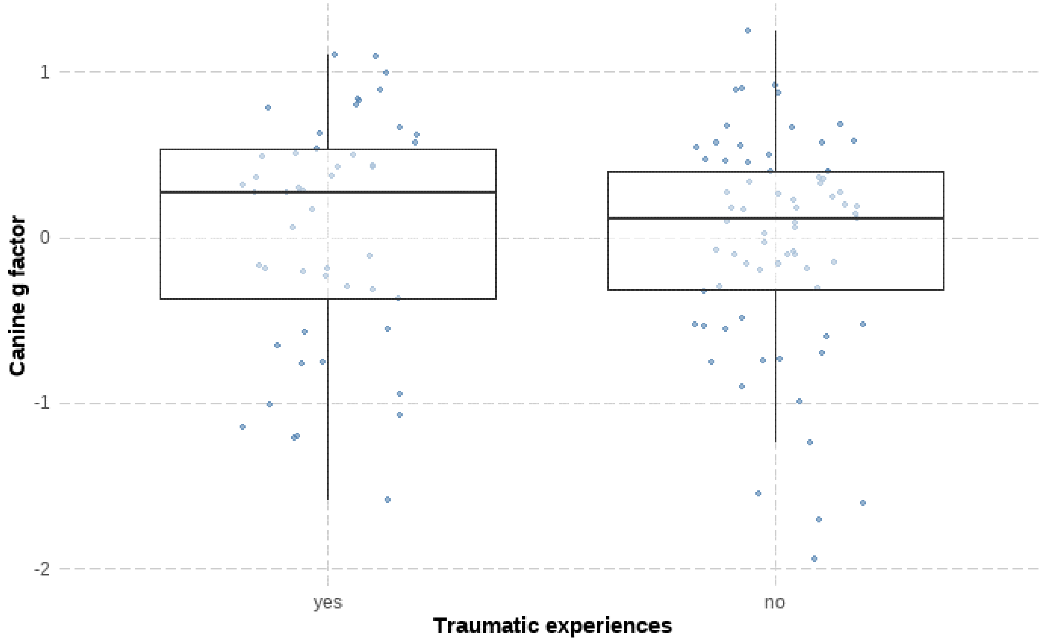


**Fig. S5. Dogs that experienced trauma had higher *canine g* factor scores than those who did not**

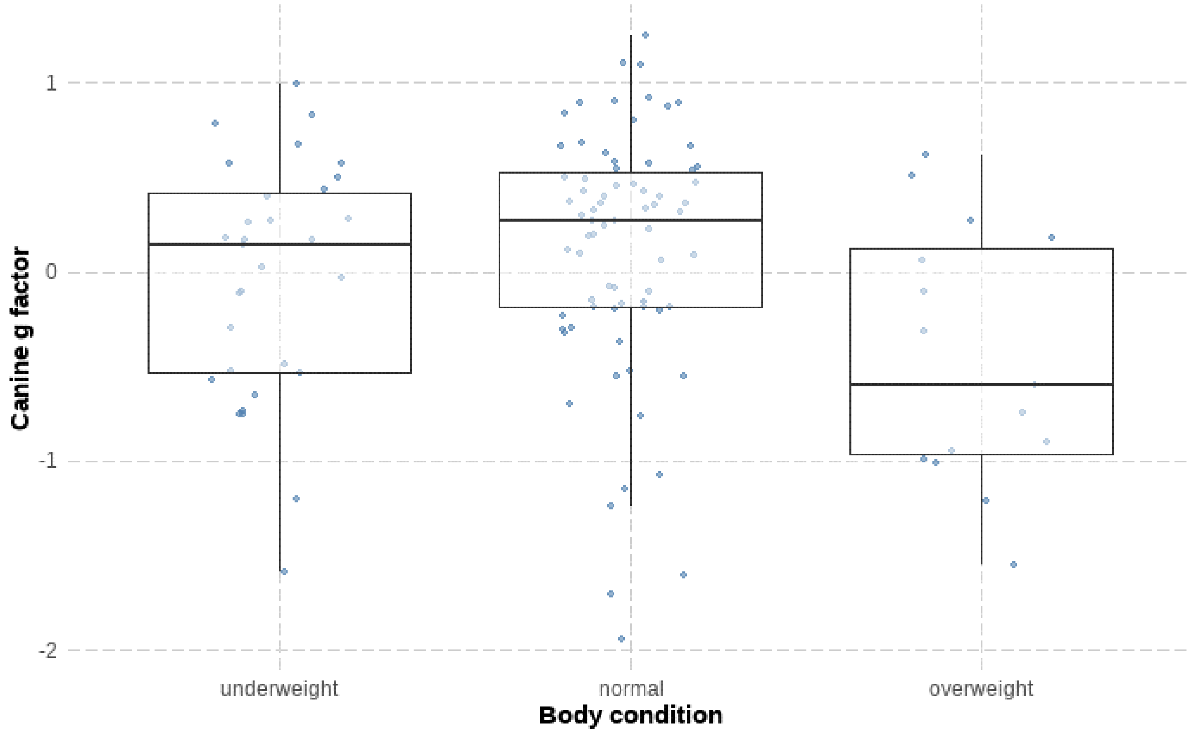


**Fig. S6. Dogs with normal body condition had higher *canine g* factor scores than the other two body condition groups**

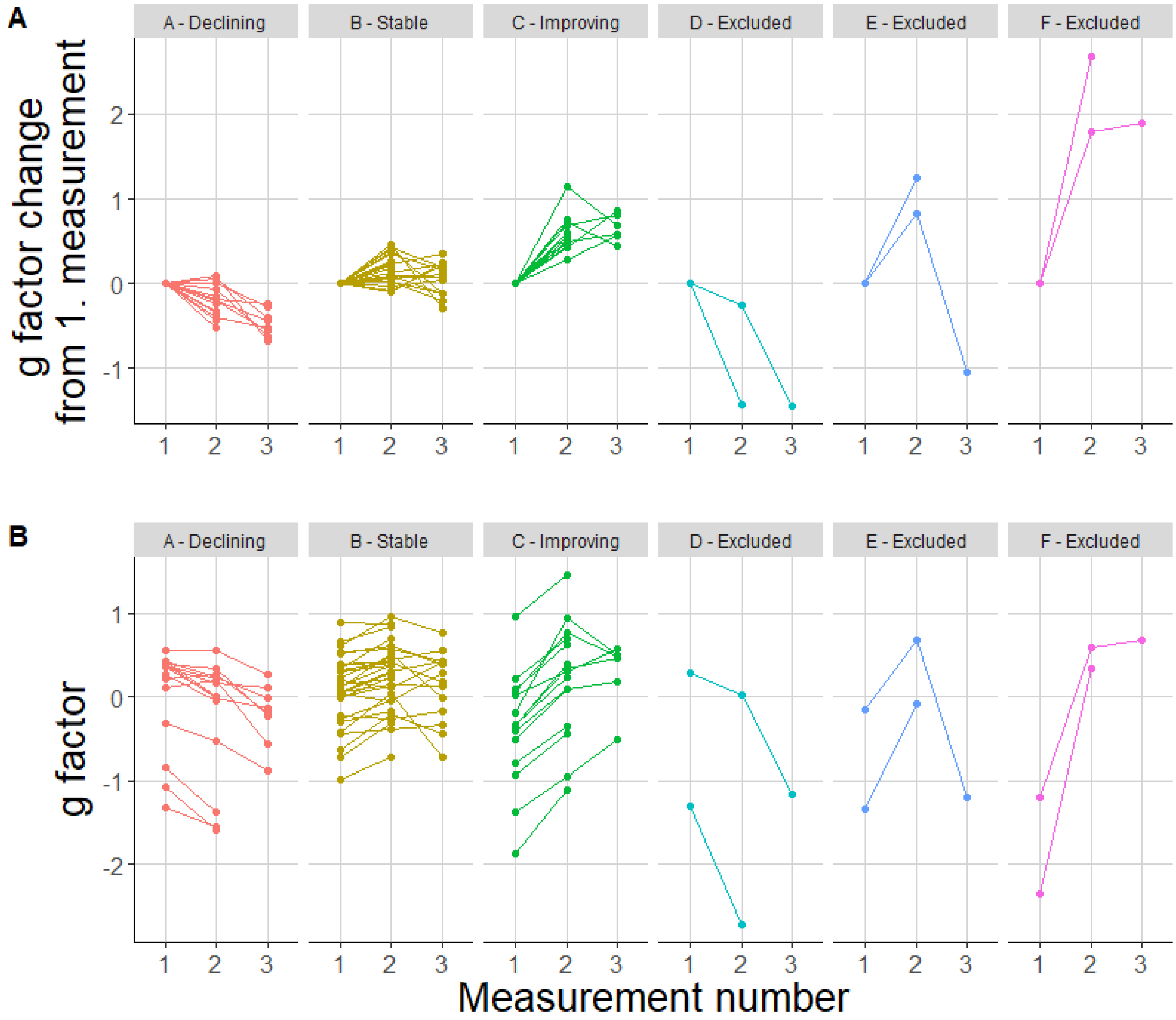


**Fig. S7. Six clusters of trajectories of dogs' *g* factor values' changes from the first measurement (A), depicted on the *g* factor scale (B).** Cluster A: N = 15, cluster B: N = 30, cluster C: N = 15, cluster D: N = 2, cluster E: N = 2, cluster F: N = 2

**Table S1. Definition and inter-observer reliability (assessed by intraclass correlation (ICC)) of the variables coded in the eleven tasks in the Cognitive battery.** E: experimenter, O: owner

| Task/Phase | Variable | Definition | ICC | p |
|---|---|---|---|---|
| COGNITIVE TASKS | | | | |
| **1. Pointing** (*80*) | | | | |
| test trials | N of correct choices | Frequency of correct choices (pot with food) (out of 6) | Reported by the E | |
| **2. Manipulative persistency** (*81*), main domain(s) assessed: Persistency, Food motivation | | | | |
| solvable | Time% look at the Kong | Duration of looking at the Kong | 0.99 | <0.001 |
| solvable | Time% touch the Kong | Duration of manipulating (touching, pawing, mouthing, nosing) the Kong | 0.96 | <0.001 |
| unsolvable | Time% look at the Kong | Duration of looking at the Kong | 0.99 | <0.001 |
| unsolvable | Time% touch the Kong | Duration of manipulating (touching, pawing, mouthing, nosing) the Kong | 0.99 | <0.001 |
| **3. Clicker game** (*82*) | | | | |
| | N of repeated behaviors | Frequency of previously rewarded behaviors repeated (except looking at E) | 0.76 | <0.001 |
| | N of novel behaviors | Frequency of new behaviors | 0.98 | <0.001 |
| | Time% repeated behaviors | Duration of performing previously rewarded behaviors (except looking at E) | 0.97 | <0.001 |
| | Time% passivity | Duration of staying still (sitting, lying, standing in one place) | 0.96 | <0.001 |
| | N of food-related behaviors | Frequency of behaviors aimed at obtaining the food (touching E's hand, the clicker, or the food bag, circling or jumping at E) | 0.42 | 0.028 |
| | N of obedience tricks | Frequency of previously learned tricks performed, e.g., sit, lay down, giving paw, turning around, etc. | 0.89 | <0.001 |
| **4. Problem solving** (*83*) | | | | |
| opaque | Latency to find food, 1st to 3rd trials | From the moment the dog starts moving until the dog reaches into the box (or maximum: 30sec) | 0.69-0.92 | <0.001 for all |
| transparent | Latency to find food, 4th to 10th trials | From the moment the dog starts moving until the dog reaches into the box (or maximum: 30sec) | 0.82-0.99 | <0.001 for all |
| all trials | N of correct first choices | Frequency of times the dog finds the food in the first choice (out of 10) | 0.70 | <0.001 |

| Task/Phase | Variable | Definition | ICC | p |
|---|---|---|---|---|
| **5. Attention** (*37*, *84*) | | | | |
| non-social | Latency to look away object | From the moment the frisbee starts moving until the dog looks away: 0: < 5 sec; 1: 5 - < 16 sec; 2: 16 - < 30 sec; 3: => 30 sec | 0.92 | <0.001 |
| non-social | Time% look at the object | 0: nearly 0%; 1:<50%; 2: =>50%; 3: nearly 100% of the time | 0.61 | 0.002 |
| social | Latency to look away human | From the moment the door opens until the dog looks away: 0: < 5 sec; 1: 5 - < 16 sec; 2: 16 - < 30 sec; 3: => 30 sec | 0.95 | <0.001 |
| **6. Training for eye contact** (*37*, *82*) | | | | |
| training | Mean eye contact latency | Mean latency of the first 15 eye contacts (from the moment the dog takes the sausage into its mouth until E clicks) (or maximum: 60 sec) | 1.00 | <0.001 |
| training | Latency to learn | If the dog passed the training criteria, the sum of the first 15 eye contact's latency; if the dog did not pass the training criteria: 300 sec | 1.00 | <0.001 |
| sustained | Sustained eye contact | Maximum duration of sustained eye contact: 0: did not pass first level; 1: 2 sec; 2: 5 sec; 3: 10 sec; 4: 20 sec; 5: 40 sec. Mean of the with and without distraction conditions | Reported by E | |
| **7. Memory** (*85*) | | | | |
| | Latency to find food, 1st to 5th trials | From the moment the dog starts moving until the dog's nose enters into the pot (or maximum: 30sec) | 0.83-1.00 | <0.001 for all |
| | N of correct first choices | Frequency of times the dog finds the food in the first choice (out of 5) | 1.00 | <0.001 |
| TESTS USED FOR VALIDATING THE *CANINE G* | | | | |
| **Exploration** (*86*) | | | | |
| on leash | Time% activity | Moving the legs: 0: nearly 0%; 1: <50%; 2: =>50%; 3: nearly 100% of the time | 0.71 | <0.001 |
| free | Time% activity | Moving the legs: 0: nearly 0%; 1: <50%; 2: =>50%; 3: nearly 100% of the time | 0.87 | <0.001 |
| free | Time% proximity to O | being < 1 m to O: 0: nearly 0%; 1: <50%; 2: =>50%; 3: nearly 100% of the time | 0.75 | <0.001 |
| free | N of objects visited | nose < 10 cm from object: 0: 0-1 objects; 1: 2-5 objects; 2: 6-10 objects; 3: 11-16 objects | 0.97 | <0.001 |

| Task/Phase | Variable | Definition | ICC | p |
|---|---|---|---|---|
| **Box rustle** (*86*) | | | | |
| | Search box, 1st to 3rd | When O investigates the box, the dog 0: does not look; 1: looks but no approach; 2: approaches <1m but no touch; 3: touches the box | 0.75-0.88 | <0.001 for all |
| | Time% following O | When O walks among the boxes, the dog follows O: 0: nearly 0%; 1: <50%; 2: =>50%; 3: nearly 100% of the time | 0.77 | <0.001 |
| **Novel object recognition** (*87*) | | | | |
| passive familiarization | N of approach any toys | Frequency of approaching any of the toys <20cm | 0.69 | <0.001 |
| passive familiarization | N of touch any toys | Frequency of touching any of the toys | 0.83 | <0.001 |
| passive familiarization | Time% proximity to any toys | Duration of being <20cm to any of the toys | 0.86 | <0.001 |
| passive familiarization | Time% touch any toys | Duration of touching any of the toys | 0.81 | <0.001 |
| test phase | N of approach known toy | Frequency of approaching the known toy <20cm | 0.65 | <0.001 |
| test phase | N of approach new toy | Frequency of approaching the new toy <20cm | 0.49 | 0.014 |
| test phase | Time% proximity to known toy | Duration of being <20cm to the known toy | 0.93 | <0.001 |
| test phase | Time% proximity to new toy | Duration of being <20cm to the new toy | 0.93 | <0.001 |
| test phase | N of touch known toy | Frequency of touching the known toy | 0.53 | 0.008 |
| test phase | N of touch new toy | Frequency of touching the new toy | 0.49 | 0.014 |
| test phase | Time% touch known toy | Duration of touching the known toy | 0.85 | <0.001 |
| test phase | Time% touch new toy | Duration of touching the new toy | 0.92 | <0.001 |
| **Reversal learning** (*53*) | | | | |
| discrimination | Number of learning trials | The number of trials required to learn the initial association between the stimuli and reward until criteria; if the dog did not pass training criteria: 51. | Reported by E | |
| reversal | Number of reversal trials | The number of trials required to learn the reversed association between the stimuli and reward; if the dog did not pass training criteria: 51. | Reported by E | |

**Table S2. Description, internal consistency and task reliability assessments of the cognitive measures obtained from the seven cognitive tasks.** A PCA was run on the coded variables for six tasks (all, except the Pointing task). The task reliability has been assessed using intraclass correlation (ICC) on N = 32 dogs

| Task/Phase | Variable | Component 1 | Component 2 |
|---|---|---|---|
| **1. Pointing** | | | |
| Test trials | N of correct choices | *no PCA* | |
| *ICC* | | *-0.060, p=0.562* | |
| **2. Manipulative persistency** | | *Persistency* | |
| solvable | Duration of looking at the Kong | **0.914** | |
| solvable | Duration of manipulating the Kong | **0.907** | |
| unsolvable | Duration of looking at the Kong | **0.916** | |
| unsolvable | Duration of manipulating the Kong | **0.915** | |
| *Explained variance (%)* | | *83.336* | |
| *Cronbach's alpha* | | *0.931* | |
| *ICC* | | *0.818, p<0.001* | |
| **3. Clicker game** | | *Flexibility* | *One-trial learning* |
| | Number of repeated behaviors | 0.371 | **0.811** |
| | Number of novel behaviors | **0.646** | 0.378 |
| | Duration of repeated behaviors | **-0.584** | **0.668** |
| | Duration of passivity | **-0.850** | 0.006 |
| | Number of food-related behaviors | **0.706** | 0.008 |
| | Number of obedience tricks | 0.058 | **0.921** |
| *Explained variance (%)* | | *42.865* | *28.148* |
| *Cronbach's alpha* | | *0.645* | *0. 728* |
| *ICC* | | *0.569, p=0.012* | *0.806, p<0.001* |
| **4. Problem solving** | | *Problem-solving success* | |
| opaque | Latency of finding the food, 1st trial | **-0.558** | |
| opaque | Latency of finding the food, 2nd trial | **-0.782** | |
| opaque | Latency of finding the food, 3rd trial | **-0.768** | |
| transp. | Latency of finding the food, 4th trial | **-0.764** | |
| transp. | Latency of finding the food, 5th trial | **-0.831** | |
| transp. | Latency of finding the food, 6th trial | **-0.794** | |
| transp. | Latency of finding the food, 7th trial | **-0.835** | |
| transp. | Latency of finding the food, 8th trial | **-0.834** | |
| transp. | Latency of finding the food, 9th trial | **-0.818** | |
| transp. | Latency of finding the food, 10th trial | **-0.705** | |
| all trials | Number of correct first choices | does not load >0.5 | |
| *Explained variance (%)* | | *59.771* | |
| *Cronbach's alpha* | | *0.923* | |
| *ICC* | | *0.549, p=0.016* | |

| Task/Phase | Variable | Component 1 | Component 2 |
|---|---|---|---|
| **5. Attention** | | *Attention to Object* | |
| non-social | Latency of looking away from the object | **0.928** | |
| non-social | Duration of looking at the object | **0.928** | |
| social | Latency of looking away from the human | does not load >0.5 | |
| *Explained variance (%)* | | *86.050* | |
| *Cronbach's alpha* | | *0.817* | |
| *ICC* | | *0.568, p=0.012* | |
| **6. Training for eye contact** | | *Associative learning* | |
| training | Mean eye contact latency | **-0.878** | |
| training | Latency to learn | **-0.936** | |
| sustained | Sustained eye contact | **0.848** | |
| *Explained variance (%)* | | *78.852* | |
| *Cronbach's alpha* | | *0.865* | |
| *ICC* | | *0.906, p<0.001* | |
| **7. Memory** | | *Memory* | |
| | Latency of finding the food, 1st trial | **-0.598** | |
| | Latency of finding the food, 2nd trial | **-0.632** | |
| | Latency of finding the food, 3rd trial | **-0.637** | |
| | Latency of finding the food, 4th trial | **-0.686** | |
| | Latency of finding the food, 5th trial | **-0.540** | |
| | Number of correct first choices | **0.823** | |
| *Explained variance (%)* | | *43.384* | |
| *Cronbach's alpha* | | *0.732* | |
| *ICC* | | *0.700, p<0.001* | |

**Table S3. Relationship between the seven cognitive components and the age of the dogs.** The results of both the linear and quadratic regressions are shown

| **Cognitive components** | **Regression** | **$R^2$** | **F** | **df1** | **df2** | **p** |
|---|---|---|---|---|---|---|
| Persistency | Linear | 0.017 | 2.213 | 1 | 126 | 0.139 |
| | Quadratic | 0.019 | 1.215 | 2 | 125 | 0.300 |
| Flexibility | Linear | 0.038 | 5.018 | 1 | 126 | 0.027 |
| | Quadratic | 0.046 | 3.028 | 2 | 125 | 0.052 |
| One-trial learning | Linear | 0.010 | 1.305 | 1 | 126 | 0.255 |
| | Quadratic | 0.024 | 1.545 | 2 | 125 | 0.217 |
| Problem solving success | Linear | 0.025 | 3.185 | 1 | 126 | 0.077 |
| | Quadratic | 0.037 | 2.415 | 2 | 125 | 0.093 |
| Attention to object | Linear | 0.065 | 8.902 | 1 | 127 | 0.003 |
| | Quadratic | 0.088 | 6.083 | 2 | 126 | 0.003 |
| Associative learning | Linear | 0.110 | 15.632 | 1 | 127 | <0.001 |
| | Quadratic | 0.111 | 7.834 | 2 | 126 | <0.001 |
| Memory | Linear | 0.135 | 19.826 | 1 | 127 | <0.001 |
| | Quadratic | 0.151 | 11.238 | 2 | 126 | <0.001 |

**Table S4. Results of the unrotated exploratory factor analysis on the raw data and the (linear) age-residuals of the test data.** Loadings > 0.3 are in bold

| | **Raw data** | | | **Age-residuals** | | |
|---|---|---|---|---|---|---|
| **Cognitive component** | *Factor 1* | *Factor 2* | *Factor 3* | *Factor 1* | *Factor 2* | *Factor 3* |
| Persistency | **0.644** | 0.000 | **-0.414** | **0.653** | 0.198 | **-0.306** |
| Flexibility | 0.127 | 0.056 | 0.182 | 0.042 | -0.037 | 0.119 |
| One-trial learning | **0.344** | -0.214 | 0.278 | **0.341** | -0.288 | 0.217 |
| Problem solving success | **0.689** | -0.274 | -0.130 | **0.711** | -0.167 | -0.180 |
| Attention to object | 0.288 | **0.690** | -0.057 | 0.188 | **0.670** | 0.205 |
| Associative learning | **0.512** | 0.111 | **0.474** | **0.432** | -0.075 | **0.536** |
| Memory | **0.564** | -0.001 | 0.019 | **0.502** | -0.007 | -0.041 |
| Eigenvalue | 2.213 | 1.123 | 1.105 | 2.062 | 1.163 | 1.067 |
| Explained variance (%) | 31.618 | 16.048 | 15.785 | 29.452 | 16.618 | 15.240 |

**Table S5. Results of the unrotated exploratory factor analysis on components that compose the *canine g*, on the raw data and the (linear) age-residuals of the test data.** Loadings > 0.3 are in bold

| Cognitive component | Raw data<br>*Canine g* | Age-residuals<br>*Canine g* |
|---|---|---|
| Persistency | **0.607** | **0.615** |
| One-trial learning | **0.363** | **0.360** |
| Problem solving success | **0.715** | **0.728** |
| Associative learning | **0.509** | **0.399** |
| Memory | **0.558** | **0.510** |
| Eigenvalue | 2.140 | 2.039 |
| Explained variance (%) | 42.806 | 40.772 |
| Cronbach's alpha | 0.647 | 0.626 |

**Table S6. Pattern matrix of the exploratory factor analysis conducted with Oblimin rotation on the raw data and the (linear) age-residuals of the test data.** Loadings > 0.3 are in bold

| | Raw data | | Age-residuals | |
|---|---|---|---|---|
| **Cognitive component** | *Individual problem solving* | *Learning* | *Individual problem solving* | *Learning* |
| Persistency | **0.747** | -0.122 | **0.733** | -0.126 |
| One-trial learning | 0.077 | **0.389** | 0.053 | **0.456** |
| Problem solving success | **0.680** | 0.105 | **0.680** | 0.115 |
| Associative learning | -0.058 | **0.751** | -0.031 | **0.631** |
| Memory | **0.438** | 0.199 | **0.461** | 0.102 |
| Eigenvalue | 2.140 | 1.031 | 2.039 | 1.077 |
| Cronbach's alpha | 0.674 | 0.477 | 0.664 | 0.460 |

**Table S7. Description and internal consistency of the components obtained from the Exploration, Box rustle, and Novel object recognition tests**

| Task/Phase | Variable | Component 1 | Component 2 |
|---|---|---|---|
| **Exploration** | | *Active* | |
| on leash | Duration of activity | **0.581** | |
| free | Duration of activity | **0.891** | |
| free | Duration of proximity to owner | **-0.823** | |
| free | Number of objects visited | **0.766** | |
| *Explained variance (%)* | | *43.104* | |
| *Cronbach's alpha* | | *0.757* | |
| **Box rustle** | | *Follow* | |
| | Search first box | **0.752** | |
| | Search second box | **0.718** | |
| | Search third box | **0.765** | |
| | Duration of following owner | **0.717** | |
| *Explained variance (%)* | | *54.499* | |
| *Cronbach's alpha* | | *0.651* | |
| **Novel object recognition** | | *Preference for novelty* | *Preference for known* |
| familiarization | Frequency of approaching any toys | **0.622** | 0.126 |
| familiarization | Frequency of touching any toys | **0.617** | 0.211 |
| test phase | Frequency of approaching new | **0.691** | 0.155 |
| test phase | Duration of being close to new | **0.890** | -0.196 |
| test phase | Frequency of touching new | **0.777** | 0.010 |
| test phase | Duration of touching new | **0.869** | -0.188 |
| familiarization | Duration of being close to any toys | 0.266 | **0.746** |
| familiarization | Duration of touching any toys | 0.193 | **0.769** |
| test phase | Frequency of approaching known | 0.326 | **0.565** |
| test phase | Duration of being close to known | -0.238 | **0.957** |
| test phase | Frequency of touching known | 0.067 | **0.721** |
| test phase | Duration of touching known | -0.261 | **0.916** |
| *Explained variance (%)* | | *24.203* | *39.833* |
| *Cronbach's alpha* | | *0.848* | *0.886* |

**Table S8. Demography, personality, and behavior-related questionnaire items**

| | |
|---|---|
| 1-8) Basic information about the dog and the owner: | 1) dog's age, 2) sex, 3) reproductive status, 4) breed (grouped later into purebreds and mixed breeds), 5) weight, 6) height at withers, 7) body condition (underweight, normal, overweight), 8) owner's age |
| 9) Previous experienced trauma (e.g. accident, surgery, lost for days, changed owners) | Yes/no |
| 10) Time spent playing with the owner per day | <1h / >1h |
| 11) Off leash activity per day | <1h / 1-3h / >3h |
| 12) Time spent alone per day | None / 1-2h / 3-8h / >8h |
| 13) Other dogs in the household | None / one / more |
| 14) Training level | Average of the number of commands the dog performs reliably (maximum 3/4/5/6/7 or more), the number of commands known by the dog according to the owner (less than 10/10 or more), number of training the dog fulfilled (0/1/2 or more), the number of training activities the dog currently attends (0/1/2 or more), the number of leisure activities (1 or less/2/3 or more) and the owner experience score (minimal experience/moderate/professional) |
| 15) Family level | Average of the number of people living in the household (1/2/3/4 or more) and the score value of the presence of kids (no/yes) in the household |
| 16) Health issues | Average of the score value of the question about the necessary regular medication (no/yes), treating the dog with vitamins/supplements (almost never/rarely/often/regularly[daily]) and the number of health problems listed by the owner (no problems/1/2/3 or more) |
| 17-21) Dog personality traits: | 17) Fearfulness, 18) Aggression towards People, 19) Activity/Excitability, 20) Responsiveness to Training, 21) Aggression towards Animals |
| 22) Owner's emotional attitude | Average score from the following items: my dog thinks like a child; I often pet and stroke my dog; my dog lets me know s/he needs me; my dog makes me laugh, and I often play with her/him; I always buy gifts for my dog during the holidays; my dog is more important to me than any other people |
| 23) Owner's doggy lifestyle | Average score from the following items: I try to read scientific dog literature; I talk a lot about dogs with friends; thanks to my dog, I have pleasant free time activities; I bring my dog with me even when it is not necessary |
| 24) Communication | Average score from the following items: my dog 'tells' me when hungry; calls me to play; calls for help; looks for my gaze; shows me things or lead me to things; empathic; follows my orientation |

| | |
|---|---|
| | when I look at something in the distance; changes its behavior when I smile at her/him; attentive when I say something to her/him; looks for contact with me; laughs when we meet someone s/he likes; follows people's conversation and tries to find out what refers to her/him; looks or goes to the direction I point to; tries to join in activities, tries to imitate what I do |
| 25) Signs of decline | Average score from the following items: my dog walks up and down or in circles; get lost at familiar places; stares with no reason; get stuck; has housebreaking problems; has problems with eating/drinking; doesn't react to her/his name, doesn't recognize when spoken to; shakes, trembles with no reason; clumsy, falls off the stairs; scared of familiar people, s/he does not recognize them; sensitive to weather fronts |
| 26) Uncontrollability | Average score from the following items: my dog disobeys the commands; actively tries to get attention; gets engaged into things and hard to get her/his attention; in frustrating situations s/he starts to have tantrums; jumps up on people |
| 27) Sociability/trainability | Average score from the following items: my dog follows me from room to room; has recognizable emotions/feelings; greets enthusiastically; remembers well earlier taught things; learns quickly new tasks; good in holding back urine or faeces |

**Table S9. AIC-based parsimonious model for Model1: Individual features**

| *Predictors* | *Estimates* | *CI* | *Statistic* | *p* |
|---|---|---|---|---|
| (Intercept) | 4.854 | 1.930 – 7.778 | 3.291 | **0.001** |
| dog's age | -0.165 | -0.480 – 0.149 | -1.042 | 0.300 |
| reproductive status [neutered] | 0.483 | -0.722 – 1.689 | 0.794 | 0.429 |
| height | -0.027 | -0.045 – -0.009 | -2.988 | **0.003** |
| body condition [normal] | 0.408 | -0.016 – 0.832 | 1.908 | 0.059 |
| body condition [overweight] | -0.378 | -1.057 – 0.300 | -1.106 | 0.271 |
| health issues | 0.966 | 0.060 – 1.872 | 2.114 | **0.037** |
| training level | -0.051 | -1.140 – 1.039 | -0.092 | 0.927 |
| trauma [no] | -1.565 | -2.702 – -0.429 | -2.730 | **0.007** |
| dog's age * reproductive status [neutered] | -0.136 | -0.289 – 0.018 | -1.748 | 0.083 |
| dog's age * health issues | -0.134 | -0.236 – -0.032 | -2.599 | **0.011** |
| dog's age * training level | 0.121 | 0.001 – 0.242 | 2.000 | **0.048** |
| dog's age * trauma [no] | 0.124 | -0.002 – 0.249 | 1.945 | 0.054 |
| Observations | 120 | | | |
| $R^2$ / $R^2$ adjusted | 0.498 / 0.442 | | | |

**Table S10. AIC-based parsimonious model for Model2: Keeping conditions**

| *Predictors* | *Estimates* | *CI* | *Statistic* | *p* |
|---|---|---|---|---|
| (Intercept) | 4.207 | 3.548 – 4.867 | 12.648 | **<0.001** |
| dog's age | -0.155 | -0.233 – -0.076 | -3.925 | **<0.001** |
| Observations | 104 | | | |
| $R^2$ / $R^2$ adjusted | 0.131 / 0.123 | | | |

**Table S11. AIC-based parsimonious model for Model3: Personality scores**

| *Predictors* | *Estimates* | *CI* | *Statistic* | *p* |
|---|---|---|---|---|
| (Intercept) | 5.032 | -0.510 – 10.573 | 1.799 | 0.075 |
| dog's age | -0.675 | -1.257 – -0.094 | -2.301 | **0.023** |
| Aggression towards People | -0.958 | -2.047 – 0.132 | -1.742 | 0.084 |
| Activity/Excitability | 0.515 | 0.100 – 0.931 | 2.456 | **0.016** |
| Responsiveness to Training | -0.358 | -1.421 – 0.704 | -0.668 | 0.506 |
| dog's age * Aggression toward People | 0.117 | -0.029 – 0.262 | 1.588 | 0.115 |
| dog's age * Responsiveness to Training | 0.097 | -0.022 – 0.217 | 1.611 | 0.110 |
| Observations | 121 | | | |
| $R^2$ / $R^2$ adjusted | 0.266 / 0.227 | | | |

**Table S12. AIC-based parsimonious model for Model4: Query scores**

| *Predictors* | *Estimates* | *CI* | *Statistic* | *p* |
|---|---|---|---|---|
| (Intercept) | 6.614 | 1.629 – 11.600 | 2.628 | **0.010** |
| dog’s age | -0.420 | -0.856 – 0.017 | -1.905 | 0.059 |
| emotional | -0.452 | -0.780 – -0.125 | -2.734 | **0.007** |
| doggy lifestyle | -0.419 | -1.384 – 0.547 | -0.859 | 0.392 |
| Signs of decline | -1.323 | -2.580 – -0.066 | -2.085 | **0.039** |
| Sociability/trainability | 0.569 | 0.148 – 0.990 | 2.678 | **0.008** |
| dog’s age * doggy lifestyle | 0.072 | -0.031 – 0.175 | 1.392 | 0.167 |
| Observations | 121 | | | |
| $R^2$ / $R^2$ adjusted | 0.271 / 0.233 | | | |

**Table S13. AIC-based parsimonious model for Combined model**

| | **F test results** | | | | | |
|---|---|---|---|---|---|---|
| | *Df* | *Sum of Squares* | *Residual Sum of Squares* | *AIC* | *F value* | *P value* |
| <none> | | | 87.397 | -4.043 | | |
| **height** | **1** | **7.926** | **95.323** | **4.374** | **9.341** | **0.0029** |
| **body condition** | **2** | **9.749** | **97.146** | **4.647** | **5.745** | **0.0044** |
| **training level** | **1** | **8.126** | **95.523** | **4.625** | **9.576** | **0.0026** |
| **trauma** | **1** | **10.366** | **97.764** | **7.407** | **12.218** | **0.0007** |
| **Activity/Excitability** | **1** | **7.655** | **95.052** | **4.032** | **9.021** | **0.0034** |
| **emotional** | **1** | **3.705** | **91.103** | **-1.061** | **4.367** | **0.0391** |
| dog age * reproductive status | *1* | *3.070* | *90.468* | *-1.900* | *3.618* | *0.0599* |
| **dog age * health issues** | **1** | **7.871** | **95.268** | **4.304** | **9.276** | **0.0029** |
| *dog age * Responsiveness to training* | *1* | *2.666* | *90.063* | *-2.437* | *3.142* | *0.0793* |
| dog age * doggy lifestyle | 1 | 2.277 | 89.677 | -2.953 | 2.687 | 0.1042 |

**Table S14. AIC-based parsimonious model for Combined model, controlled for multiple comparisons**

| | F test results | | | | | |
|---|---|---|---|---|---|---|
| | *Df* | *Sum of Squares* | *Residual Sum of Squares* | *AIC* | *F value* | *P value* |
| <none> | | | 101.14 | 1.478 | | |
| **height** | **1** | **6.155** | **107.29** | **6.567** | **6.634** | **0.0113** |
| **body condition** | **2** | **8.076** | **109.21** | **6.697** | **4.352** | **0.0152** |
| **training level** | **1** | **12.914** | **114.05** | **13.898** | **13.918** | **0.0003** |
| **trauma** | **1** | **8.547** | **109.68** | **9.213** | **9.212** | **0.0030** |
| **Activity/Excitability** | **1** | **13.849** | **114.98** | **14.878** | **14.926** | **0.0002** |
| **emotional** | **1** | **4.066** | **105.20** | **4.209** | **4.384** | **0.0386** |
| **dog_age * health issues** | **1** | **10.031** | **111.17** | **10.825** | **10.810** | **0.0014** |

**Table S15. Details of the AIC-based parsimonious model for Combined model, controlled for multiple comparisons**

| | Canine *g* factor | | | |
|---|---|---|---|---|
| *Predictors* | *Estimates* | *CI* | *Statistic* | *p* |
| (Intercept) | 3.110 | 2.694 – 3.526 | 14.804 | <0.001 |
| dog's age | -0.360 | -0.554 – -0.166 | -3.685 | <0.001 |
| height | -0.234 | -0.413 – -0.054 | -2.576 | 0.011 |
| body condition [normal] | 0.493 | 0.076 – 0.910 | 2.344 | 0.021 |
| body condition [overweight] | -0.144 | -0.786 – 0.497 | -0.445 | 0.657 |
| health issues | -0.048 | -0.238 – 0.141 | -0.507 | 0.613 |
| training level | 0.377 | 0.177 – 0.577 | 3.731 | <0.001 |
| trauma [no] | -0.566 | -0.936 – -0.196 | -3.035 | 0.003 |
| Activity/Excitability | 0.402 | 0.196 – 1.608 | 3.863 | <0.001 |
| emotional | -0.197 | -0.384 – -0.011 | -2.094 | 0.039 |
| dog's age * health issues | -0.341 | -0.546 – -0.135 | -3.288 | 0.001 |
| Observations | 120 | | | |
| R2 / R2 adjusted | 0.508 / 0.463 | | | |

**Table S16. The results of the classification based on the trajectories of *g* score changes**

| Clusters (dogs' ratio) | g score change (mean ± SD) | |
|---|---|---|
| | *from 1. to 2. measurement* | *from 2. to 3. measurement* |
| A (22.7%) | -0.23 ± 0.20 | -0.36 ± 0.24 |
| B (45.5%) | 0.17 ± 0.16 | -0.09 ± 0.30 |
| C (22.7%) | 0.59 ± 0.20 | 0.03 ± 0.34 |
| D (3.0%) | -0.84 ± 0.82 | -1.20 |
| E (3.0%) | 1.00 ± 0.30 | -1.90 |
| F (3.0%) | 2.20 ± 0.63 | 0.10 |

**Table S17. Result of bottom-up model selection for identifying the factors influencing the cluster membership in the longitudinal analysis**

| Model comparisons | **One-variable model** compared to null model (AIC = 129) | | **Two-variables model** compared to one-variable model (AIC = 126) | |
|---|---|---|---|---|
| Variables | AIC | p | AIC | p |
| *g* score at the first measurement | **126** | **0.035 *** | Already in the model | |
| dog's age | 131 | 0.447 | 129 | 0.582 |
| sex | 129 | 0.171 | 126 | 0.166 |
| breed | 133 | 0.967 | 130 | 0.945 |
| weight | 132 | 0.673 | 129 | 0.627 |
| height | 132 | 0.629 | 129 | 0.473 |
| body condition | 134 | 0.583 | 131 | 0.569 |
| previously experienced trauma | 132 | 0.730 | 128 | 0.308 |
| time spent playing | 131 | 0.442 | 129 | 0.645 |
| off-leash activity | 135 | 0.799 | 132 | 0.738 |
| time spent alone | 131 | 0.149 | 126 | 0.067 |
| other dogs in the household | 132 | 0.344 | 129 | 0.270 |
| Training level | 130 | 0.198 | 127 | 0.262 |
| Family level | 131 | 0.473 | 129 | 0.527 |
| Health issues | 131 | 0.402 | 129 | 0.515 |
| Fearfulness | 131 | 0.488 | 128 | 0.420 |
| Aggression towards People | 133 | 0.900 | 130 | 0.847 |
| Activity/Excitability | 132 | 0.615 | 129 | 0.523 |
| Responsiveness to Training | 131 | 0.458 | 129 | 0.589 |
| Aggression towards Animals | 130 | 0.240 | 128 | 0.280 |
| Owner's emotional attitude | 131 | 0.370 | 126 | 0.103 |
| Owner's doggy lifestyle | 132 | 0.540 | 129 | 0.565 |

**Table S18. Differences between the different clusters**

| | **Clusters based on *g* score change** | | |
|---|---|---|---|
| | *A - Declining* | *B - Stable* | *C - Improving* |
| **Initial *g* factor score** (mean ± SD) | **0.02 ± 0.61** | **0.07 ± 0.43** | **-0.39 ± 0.68** |
| vs A - Declining | - | - | - |
| vs B - Stable | - | - | ß = 1.43 ± 0.61, Z = 2.35, p = 0.019 |
| vs C - Improving | - | ß = -1.43 ± 0.61, Z = -2.35, p = 0.019 | - |

**Movie S1. (separate file)**

Protocol video of the cognitive tasks assessing the canine g factor.

**Data S1. (separate file)**

Data of all test occasions, g factor scores of different analyses, basic, environment-, and behavior-related information of dogs, dog personality traits scores, and owner's attitude information.
